# Piggybacking on niche-adaptation reduces the cost of multidrug resistance plasmids

**DOI:** 10.1101/2020.10.23.351932

**Authors:** Julia Kloos, João A. Gama, Joachim Hegstad, Ørjan Samuelsen, Pål J. Johnsen

## Abstract

The persistence of plasmids in bacterial populations represents a puzzling evolutionary problem with serious clinical implications due to their role in the ongoing antibiotic resistance crisis. Recently, major advancements have been made towards resolving this “plasmid paradox” but mainly in a non-clinical context. Here we propose an additional explanation for the maintenance of multidrug resistance (MDR) plasmids in clinical *Escherichia coli* strains. After co-evolving two MDR plasmids encoding last resort carbapenem resistance with an extraintestinal pathogenic *E. coli* strain, we observed that chromosomal media adaptive mutations in the global regulatory systems CCR (Carbon Catabolite Repression) and ArcAB (Aerobic Respiration Control) pleiotropically mitigated the costs of both plasmids. Mechanistically, cost reductions were due to a net downregulation of plasmid gene expression. Our results suggest that global chromosomal transcriptional re-wiring during bacterial niche-adaptation may facilitate plasmid maintenance.

## Introduction

Plasmids are self-replicating extrachromosomal elements that decrease bacterial fitness due to the requirement of host functions for their own replication and spread [1, 2]. These genetic elements play a key role in the evolution and spread of antibiotic resistance determinants in bacterial populations world-wide [3, 4]. This is particularly true for nosocomial pathogens in the family *Enterobacteriaceae* including *Escherichia coli* and *Klebsiella pneumoniae* where resistance determinants of high clinical relevance such as carbapenemases and extended-spectrum β-lactamases are frequently encoded on plasmids [5, 6].

From an evolutionary perspective, persistence of plasmids in bacterial populations has for a long time been a conundrum often referred to as the “plasmid-paradox” [7]. This paradox can be resolved in at least five different ways. First, maintenance can be ensured by positive selection for plasmid encoded traits [8–10]. But, if too beneficial, positively selected traits will be captured by the chromosome rendering the plasmid obsolete and consequently lost, as demonstrated theoretically [11] and experimentally [12]. Second, mathematical models predict that high rates of horizontal plasmid transfer can counteract segregational plasmid loss and the competitive disadvantage of plasmid-carriers [13]. *In vitro* studies report that conditions exist where conjugation frequencies are indeed extremely high [14]. It is however generally accepted that conjugation is a costly process [2] and evolution towards increased conjugation rates does not constitute a general solution of the paradox [15–17]. Third, transmissible plasmids under purifying selection may “escape” their host and enter less hostile environments. This has been termed cross-ecotype transfer [11]. Fourth, plasmid stability can evolve through improved replication control and the acquisition of addiction mechanisms [18, 19]. Fifth, and perhaps most prominent, negative effects on host fitness can be mitigated through compensatory evolution [2]. Fitness compensating mutations have been demonstrated to occur both in the presence and absence of selective agents and identified on bacterial chromosomes [10, 20, 21], on plasmids [17, 22, 23], or both [16, 24, 25].

The last ten years have brought significant advancements in the understanding of plasmid-host evolutionary dynamics. However, it is not clear how the different solutions to the plasmid paradox as listed above are relevant for clinical strains and plasmids since the majority of published work has focused on emblematic laboratory strains and/or environmental bacteria. In this report we asked if and how two clinical plasmids encoding the VIM-1 and NDM-1 carbapenemases affected fitness of an *E. coli* strain isolated from a patient, before and after experimental evolution. We observed striking parallel evolution of the CCR (Carbon Catabolite Repression) and ArcAB (Aerobic Respiration Control) regulatory systems in the chromosomes of both plasmid-containing and plasmid-free populations resulting in adaptation to the experimental conditions. No apparent compensatory mutations were identified across evolved populations and the plasmid sequences were largely unchanged. Yet, the initial plasmid costs were ameliorated in the co-evolved cultures. We demonstrate that fitness amelioration resulted from “piggybacking” on the clinical strains’ adaptation to a new niche, suggesting a novel solution to the plasmid paradox.

## Results

### Plasmid acquisition moderately reduces fitness in a clinical *E. coli* host strain

To mimic the acquisition of plasmid-mediated resistance to a last resort antibiotic, we transferred each of the two carbapenemase-producing clinical plasmids pG06-VIM-1 from *K. pneumoniae* (*bla*_VIM-1_) [26] and pK71-77-1-NDM from *E. coli* (*bla*_NDM-1_) [27] into an Extraintestinal Pathogenic *E. coli* sequence type (ST) 537 (strain ExPEC [28]) (Supplementary Table 1). pG06-VIM-1 is non-conjugative [29] while pK71-77-1-NDM is conjugative [30]. Plasmid transfer resulted in strains ExPEC+VIM and ExPEC+NDM, both otherwise isogenic to strain ExPEC (Figure 1a and Supplementary Table 1).

**Fig. 1.**
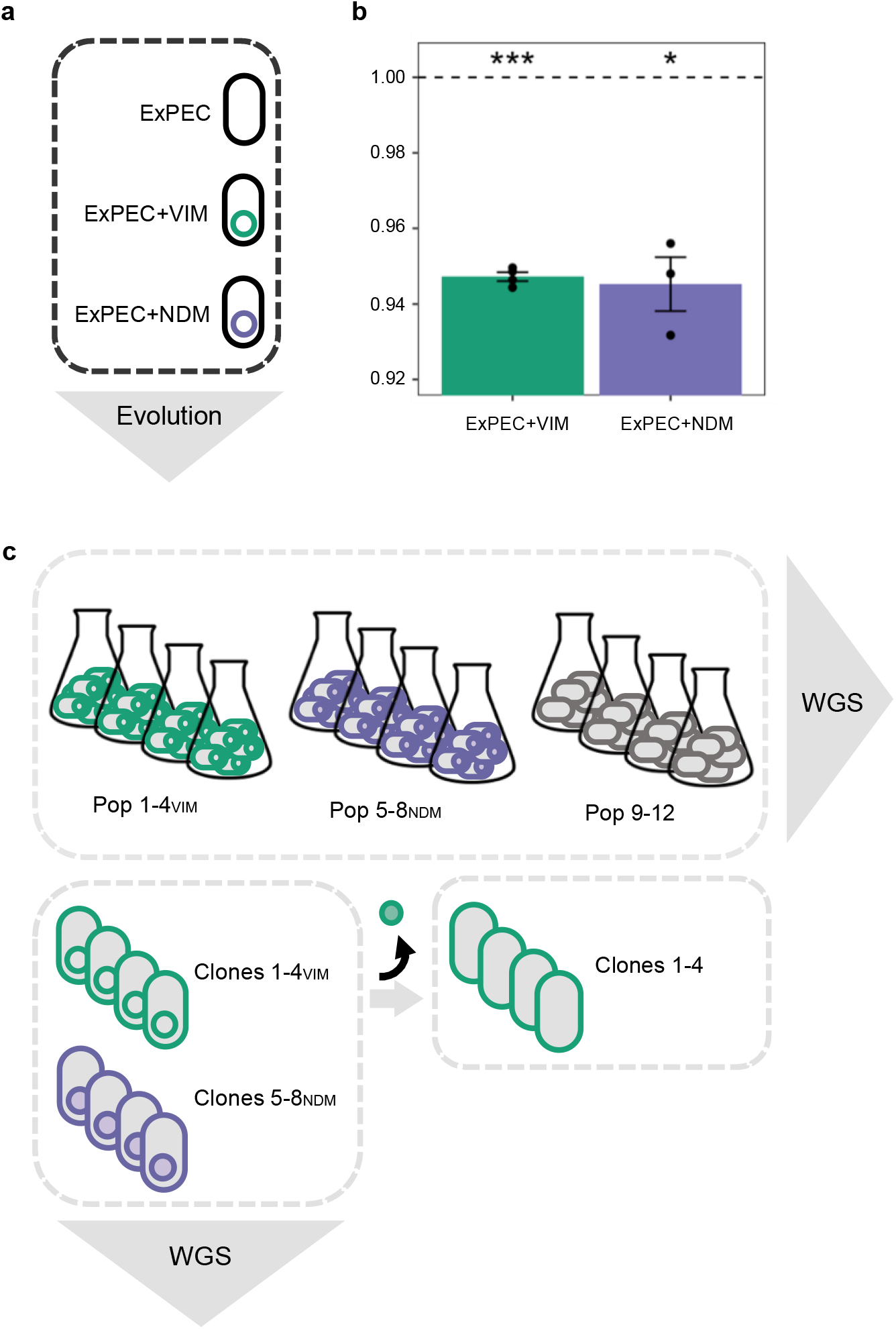
Fitness effect of plasmid acquisition and experimental procedures. **a** An ExPEC strain (black) acquired each of the two MDR plasmids pG06-VIM-1 (green; 53 kB; IncR; [29]) and pK71-77-1-NDM (purple; 145 kB; IncC; [30]) of clinical origin encoding the carbapenemases VIM-1 and NDM-1, respectively. **b** Initial fitness costs of newly transferred plasmids in strains ExPEC+VIM and ExPEC+NDM (n = 4 and 3, respectively). Significant plasmid costs are indicated by asterisks (*P* = * < 0.05, ** < 0.01, *** < 0.001; one-sample *t*-test, two-sided). Error bars indicate ± s.e.m. **c** Experimental evolution in absence of selective pressure (~300 generations) resulted in plasmid-carrying (Pop 1-4_VIM_ and Pop 5-8_NDM_) and plasmid-free (Pop 9-12) populations which were subjected to whole-genome sequencing (WGS). Representative clones per plasmid-carrying evolved population (Clones 1-4_VIM_ and Clones 5-8_NDM_) were sequenced and segregants without evolved pG06-VIM-1 (filled green circle) were generated for subsequent competition experiments (Clones 1-4).

We measured the cost of the newly introduced clinical plasmids in head-to-head competition experiments lasting ~40 generations. Acquisition of either pG06-VIM-1 or pK71-77-1-NDM affected host fitness similarly, resulting in moderate but significant costs of 5.3% and 5.5%, respectively (one-sample *t*-test, two-sided: ExPEC+VIM: *w* = 0.947 ± 0.002, *P* < 0.001; ExPEC+NDM: *w* = 0.945 ± 0.012, *P* = 0.017) (Figure 1b and Supplementary Table 6).

### Strong parallel evolution in global *E. coli* regulators occurs independently of plasmid-carriage

Four replicate lineages of the plasmid-containing strains ExPEC+VIM and ExPEC+NDM as well as the plasmid-free strain ExPEC were serially transferred for ~300 generations resulting in 12 evolved populations (Pop 1-4_VIM_, Pop 5-8_NDM_ and Pop 9-12; Figure 1c and Supplementary Table 1). We deep sequenced the evolved populations to identify putative mutations mitigating the fitness costs of plasmid carriage.

At the population level, no changes were identified in the evolved plasmid sequences except in Pop 5_NDM_ harboring a deletion in the evolved pK71-77-1-NDM (Supplementary information section IIIa, Supplementary Table 3 and Supplementary Figure 1). However, all 12 evolved lineages revealed patterns of extensive parallel evolution in chromosomal genes that are directly or indirectly linked to the CCR and the ArcAB regulatory systems of *E. coli* (Figure 2a). In total, 68 different mutations were identified in genes *cpdA* (3’,5’-**c**yclic adenosine monophosphate (cAMP) **p**hospho**d**iesterase), *crp* (**c**AMP **r**eceptor **p**rotein; DNA-binding transcriptional regulator), *arcA* (**a**erobic **r**espiration **c**ontrol protein; DNA-binding transcriptional regulator) and *arcB* (**a**erobic **r**espiration **c**ontrol sensor protein; histidine kinase). Evolved lineages had on average acquired eight variations in these genes ranging from three (Pop 5_NDM_) to 18 (Pop 6_NDM_) different mutations for individual populations (Supplementary Figure 2). Our data revealed 25, 12, 23 and eight unique mutations in *arcA* (717 bp), *arcB* (2337 bp), *cpdA* (828 bp) *and crp* (633 bp), respectively (Supplementary Figure 2). Among these unique mutations in the respective target genes, 12, one, three and three were found repeatedly across more than one evolved population. The majority of mutations in these genes were non-synonymous single nucleotide exchanges leading to amino acid substitutions (88%). Furthermore, Pop 2_VIM_ acquired mutations upstream and in the open reading frame of *cyaA* (adenylate cyclase; cAMP synthesis). For a detailed list of mutations identified across evolved populations, including small indels, as well as mutations found in single populations, see Supplementary Table 4. Whereas *cpdA* and *arcA* were mutation targets in all 12 populations, *crp* and *arcB* were identified in ten and four populations, respectively (Figure 2a and Supplementary Figure 2). Surprisingly, the mutation profiles were not different in populations that co-evolved with any of the plasmids compared to the plasmid-free control populations, strongly suggesting that the observed mutational changes were not plasmid-specific. Genes of the CCR and ArcAB systems are indeed frequently reported as mutational targets for adaptive responses to the experimental growth conditions occurring during laboratory evolution experiments [31, 32].

**Fig. 2.**
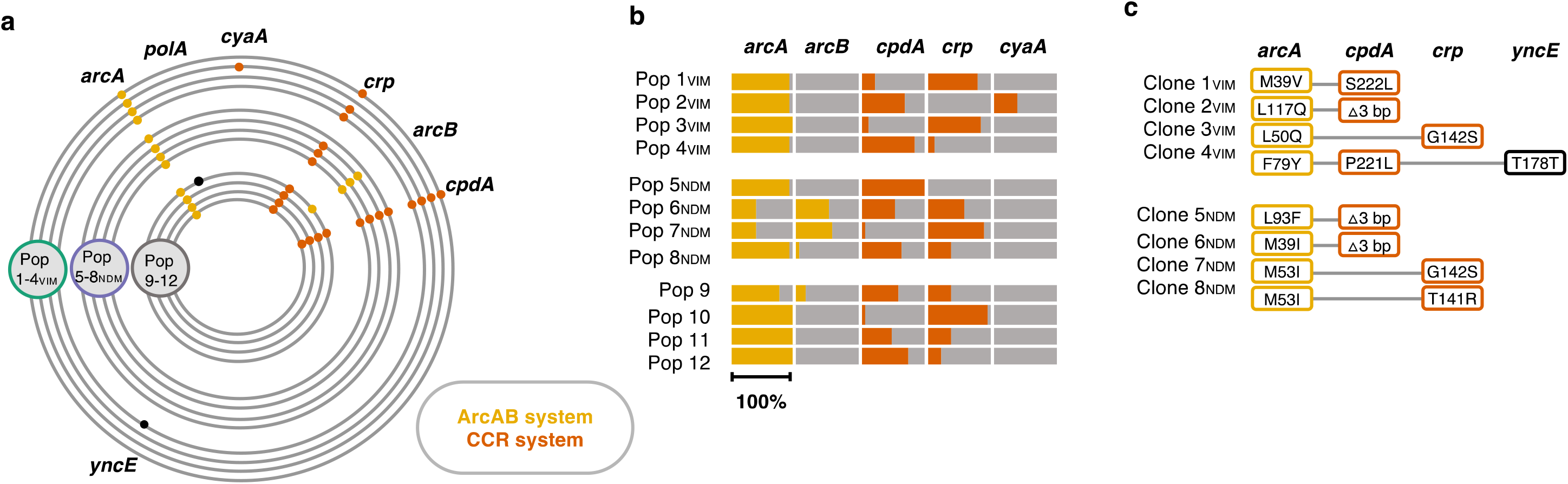
Identified mutations in the ArcAB and CCR regulatory systems. **a** Chromosomal mutations after ~300 generations of experimental evolution. Plasmid-carrying (Pop 1-4_VIM_ = green; Pop 5-8_NDM_ = purple) and plasmid-free (Pop 9-12 = grey) populations had acquired chromosomal mutations in genes associated to the ArcAB (yellow) and CCR (orange) regulatory system. Black indicates otherwise mutated genes in single evolved populations. No point mutations were identified in plasmid sequences. See also Supplementary information section IIIa and Supplementary Table 4. **b** Total frequency of all mutations targeting the same gene within single evolved populations for genes linked to ArcAB and CCR regulatory systems (entire bar length = 100%). **c** Chromosomal mutations identified in co-evolved, plasmid-carrying, whole-genome sequenced Clones 1-4_VIM_ and Clones 5-8_NDM_; ‘Δ3 bp’ in *cpdA* = *cpdA*.Δ3.bp488‑490.

### Mutations in CCR and ArcAB regulatory systems pleiotropically mitigate the cost of pG06-VIM-1 and pK71-77-1-NDM carriage

Immediate acquisitions of pG06-VIM-1 and pK71-77-1-NDM reduced host fitness significantly (Figure 1b). We have previously demonstrated complete retention of the same plasmids following experimental evolution under the same antibiotic-free conditions [29, 30]. Since one of the plasmids was non-conjugative, we assumed that fitness amelioration by compensatory adaptation was the most likely route for the plasmids to persist in evolved populations. However, the sequencing data presented above revealed no apparent compensatory mutations. Therefore, we hypothesized that adaptation to the growth conditions could have pleiotropic effects on the costs of plasmid carriage.

To test this hypothesis, we isolated a single clone from each evolved plasmid-carrying population (Pop 1-4_VIM_ and Pop 5-8_NDM_) with mutations in both regulatory systems, CCR and ArcAB, since population sequencing data suggested that both systems were affected simultaneously (Figures 2a and 2b). In the selected Clones 1-4_VIM_ and Clones 5-8_NDM_, Sanger and Illumina sequencing confirmed the presence of mutations as expected from population sequencing results and no further chromosomal or plasmid-located point mutations (Figure 1c and 2c; Supplementary information section I and IIIb, and Supplementary Table 1).

Here, we also identified large deletions in the evolved pK71-77-1-NDM for Clone 5_NDM_ and Clone 7_NDM_ (~8.8 kb and ~58.9 kb, respectively; Supplementary information section IIIb, Supplementary Table 3, Supplementary Figure 1), and susceptibility testing by disc diffusion phenotypically confirmed the deletions involving antibiotic resistance genes (Supplementary information section VI, Supplementary Table 8). The plasmid copy-number for pG06-VIM-1 and pK71-77-1-NDM before and after experimental evolution was unchanged (0.9-1.5 copies in all sequenced plasmid-carrying clones; Supplementary information section IIIb; Supplementary Table 5). Next, we attempted to generate a set of plasmid-free segregants of Clones 1-4_VIM_ and Clones 5-8_NDM_ to use in competition experiments. Plasmid curing was successful for pG06-VIM-1 resulting in Clones 1-4, but not for pK71-77-1-NDM (Figure 1c).

The costs of pG06-VIM-1-carriage in co-evolved Clones 1-4_VIM_ were assessed in head-to-head competitions with the respective plasmid-free isogenic strains (Clones 1-4) over ~40 generations. Our data show that the initial costs were significantly ameliorated to ≤ 1% in all four evolved backgrounds irrespective of the combination of chromosomal mutations in these clones (one-sample *t*-test, two-sided: Clone 1_VIM_: 0.7% or *w* = 0.993 ± 0.002, *P* = 0.017; Clone 2_VIM_: 0.4% or *w* = 0.996 ± 0.001, *P* = 0.016; Clone 3_VIM_: 0.9% or *w* = 0.991 ± 0.0003, *P* = 0.001 and Clone 4_VIM_: 0.6% or *w* = 0.994 ± 0.002, *P* = 0.056; one-way ANOVA assuming equal variances, df = 4, *P* < 0.001, followed by Dunnett’s test: *P* < 0.001; Figure 3a and Supplementary Table 6). Illumina sequencing confirmed that the plasmid sequences in Clones 1-4_VIM_ were unchanged after evolution suggesting that the chromosomal mutations were responsible for the fitness mitigation. To further test this we introduced the ancestral pG06-VIM-1 into Clone 2 and Clone 3 carrying mutations in *arcA/cpdA* and *arcA/crp*, respectively, resulting in Clone 2+VIM and Clone 3+VIM (Figure 3b). Competition experiments with the isogenic, plasmid-free genetic backgrounds revealed a significant fitness increase compared to the original plasmid-host combination and an amelioration of the initial cost of harboring pG06-VIM-1 to 1.3% and 1%, respectively (one-sample *t*-test, two-sided: Clone 2+VIM: *w* = 0.987 ± 0.004, *P* = 0.026; Clone 3+VIM: *w* = 0.990 ± 0.001, *P* = 0.002; one-way ANOVA assuming equal variances, df = 2, *P* < 0.001, followed by Dunnett’s test: *P* < 0.001; Figure 3b and Supplementary Table 6).

**Fig. 3.**
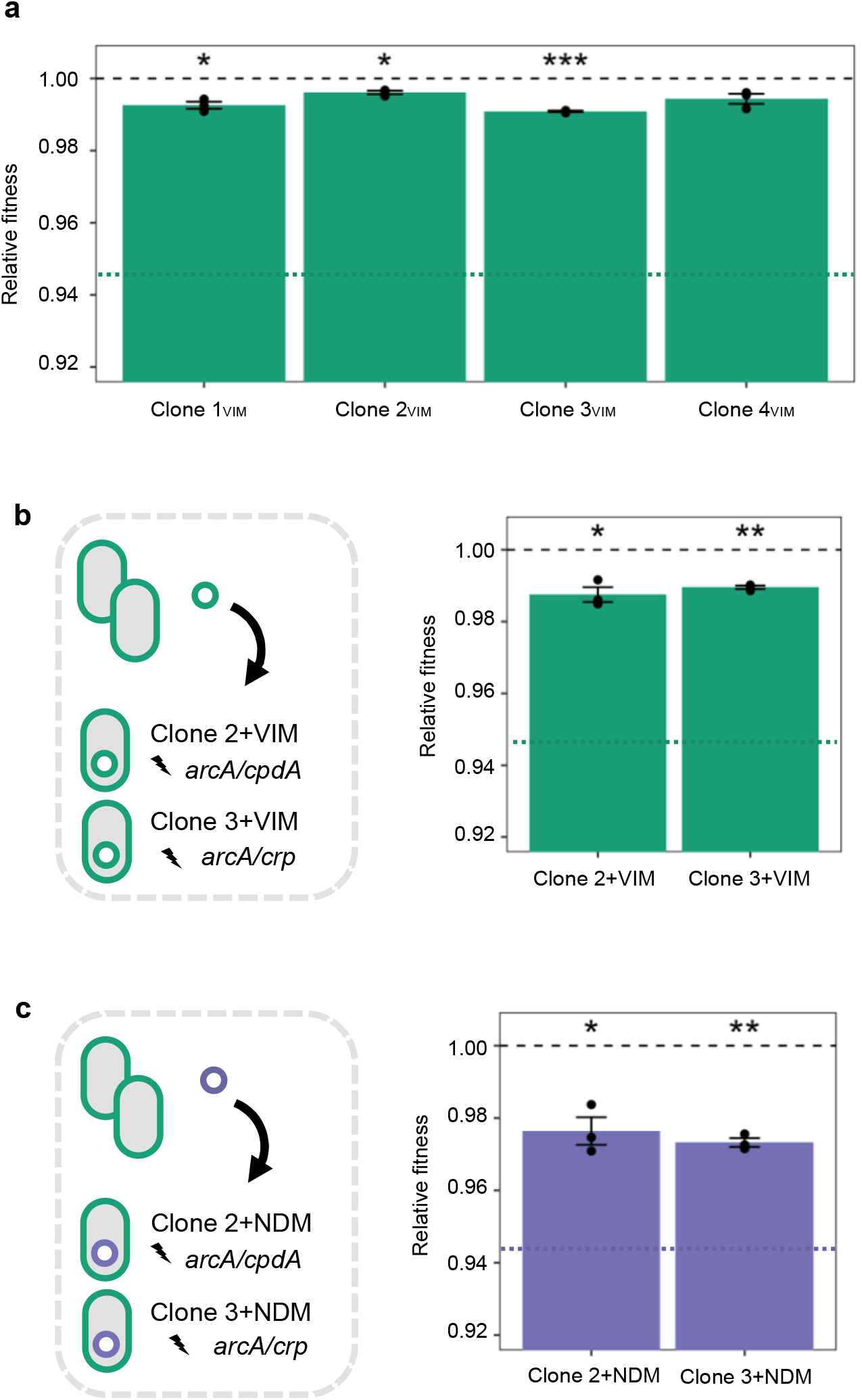
Fitness costs of evolved and ancestral plasmids in adapted backgrounds. **a** Fitness cost of co-evolved pG06-VIM-1-carrying clones (n = 3). Fitness of the ancestral strain ExPEC+VIM is indicated by a dotted green line. **b** Fitness cost of ancestral pG06-VIM-1 re-introduced into evolved Clone 2 and Clone 3. Fitness of ancestral strain ExPEC+VIM is indicated by a dotted green line. **c** Fitness cost of ancestral pK71-77-1-NDM introduced into evolved Clone 2 and Clone 3 (n = 3). Fitness of the ancestral strain ExPEC+NDM is indicated by a dotted purple line. Significant plasmid costs are indicated by asterisks (*P* = * < 0.05, ** < 0.01, *** < 0.001; one-sample *t*-test, two-sided). Error bars indicate ± s.e.m.

Similarly, the initial fitness cost of 5.5% conferred by the ancestral pK71-77-1-NDM in strain ExPEC+NDM was significantly decreased in pG06-VIM-1-free segregants carrying this plasmid (one-sample *t*-test, two-sided: Clone 2+NDM: 2.4% or *w* = 0.976 ± 0.007; *P* = 0.025; Clone 3+NDM: 2.7% or *w* = 0.973 ± 0.002; *P* = 0.002; one-way ANOVA assuming equal variances, df = 2, *P* = 0.006, followed by Dunnett’s test: *P* = 0.006 and *P* = 0.010, respectively; Figure 3c and Supplementary Table 6). Note that we did not obtain a plasmid-free background of Clones 5-8_NDM_ to test the fitness of pK71-77-1-NDM-co-evolved strains. Albeit the results from competition experiments with Clone 2+NDM and Clone 3+NDM carrying the ancestral pK71-77-1-NDM strongly suggest that the chromosomal mutations are responsible for partial fitness amelioration, we acknowledge that the deletions in evolved plasmids of Clone 5_NDM_ and Clone 7_NDM_ could result in further fitness improvements as demonstrated by Porse *et al*. [17].

Taken together, our data indicate clearly that the different mutations identified in the CCR and ArcAB regulatory systems ameliorate the initial costs of the two clinical plasmids resulting in low, if any, cost of plasmid carriage. We controlled for possible effects of plasmid loss (~40 generations) or pK71-77-1-NDM-conjugation (12 hours) in additional experiments (Methods and Supplementary information section IV and V, respectively). Both plasmids were stable (Supplementary Table 7) and the conjugation frequencies for pK71-77-1-NDM in strain ExPEC+NDM, Clone 2+NDM and Clone 3+NDM were too low to significantly affect the outcome of the competition experiments (Supplementary Figure 3).

### Plasmid cost mitigation is linked specifically to the CCR system

To investigate the individual roles of the CCR and ArcAB systems on plasmid cost mitigation we measured plasmid costs in deletion mutants for *arcA, cpdA* and *crp* – the targeted loci for adaptation in our co-evolved clones. Unfortunately, genetic modifications using clinical strains are notoriously difficult and for these experiments we used deletion mutants of *E. coli* (K-12 derivatives) from the Keio collection [33]. We introduced the ancestral pG06-VIM-1 into the individual deletion strains as well as the Keio parent strain [34] by electroporation, resulting in strains BW25113+VIM, BWΔ*cpdA*+VIM, BWΔ*arcA*+VIM and BWΔ*crp*+VIM (Supplementary information section I and Supplementary Table 1). We measured fitness of plasmid-carrying strains relative to their plasmid-free counterparts in head-to-head competitions as described above. As a general observation, pG06-VIM-1 was less costly in BW25113 than in the clinical isolate (one-sample *t*-test, two-sided: 2.3% or *w* = 0.977 ± 0.005, *P* = 0.016; Figure 4a and Supplementary Table 6). While deletion of *arcA* and *crp* had no significant effect on the fitness burden imposed by pG06-VIM-1 compared to BW25113+VIM, we measured a significant fitness improvement of the pG06-VIM-1-carrying Δ*cpdA* mutant (one-way ANOVA not assuming equal variances, df = 3, *P* = 0.004 followed by Dunnett’s test: BWΔ*arcA*+VIM, *P* = 0.993; BWΔ*crp*+VIM, *P* = 0.051; BWΔ*cpdA*+VIM, *P* = 0.001; Figure 4a and Supplementary Table 6) and a reduction of plasmid cost to 0.4% (one-sample *t*-test, two-sided: BWΔ*arcA*+VIM: 2.2% or *w* = 0.978 ± 0.007, *P* = 0.007; BWΔ*crp*+VIM: no cost or *w* = 1.006 ± 0.020, *P* = 0.534; BWΔ*cpdA*+VIM: *w* = 0.996 ± 0.002, *P* = 0.034; Figure 4a and Supplementary Table 6). These data strongly suggest that beyond the known effect on adaptation to growth conditions, *cpdA* mutations identified in our study pleiotropically mitigated plasmid costs. We could however not obtain the same level of consistency across biological replicates in competitions using the Δ*crp* mutant even though CpdA and CRP are tightly linked in the CCR regulatory system [35, 36]. Based on the observation that pG06-VIM-1 no longer imposes a cost in the Δ*crp* mutant, as confirmed by the one-sample *t*-test, we argue that *crp* is also involved in fitness mitigation despite the relatively high variance that renders the Dunnett’s test (borderline) not significant.

**Fig. 4.**
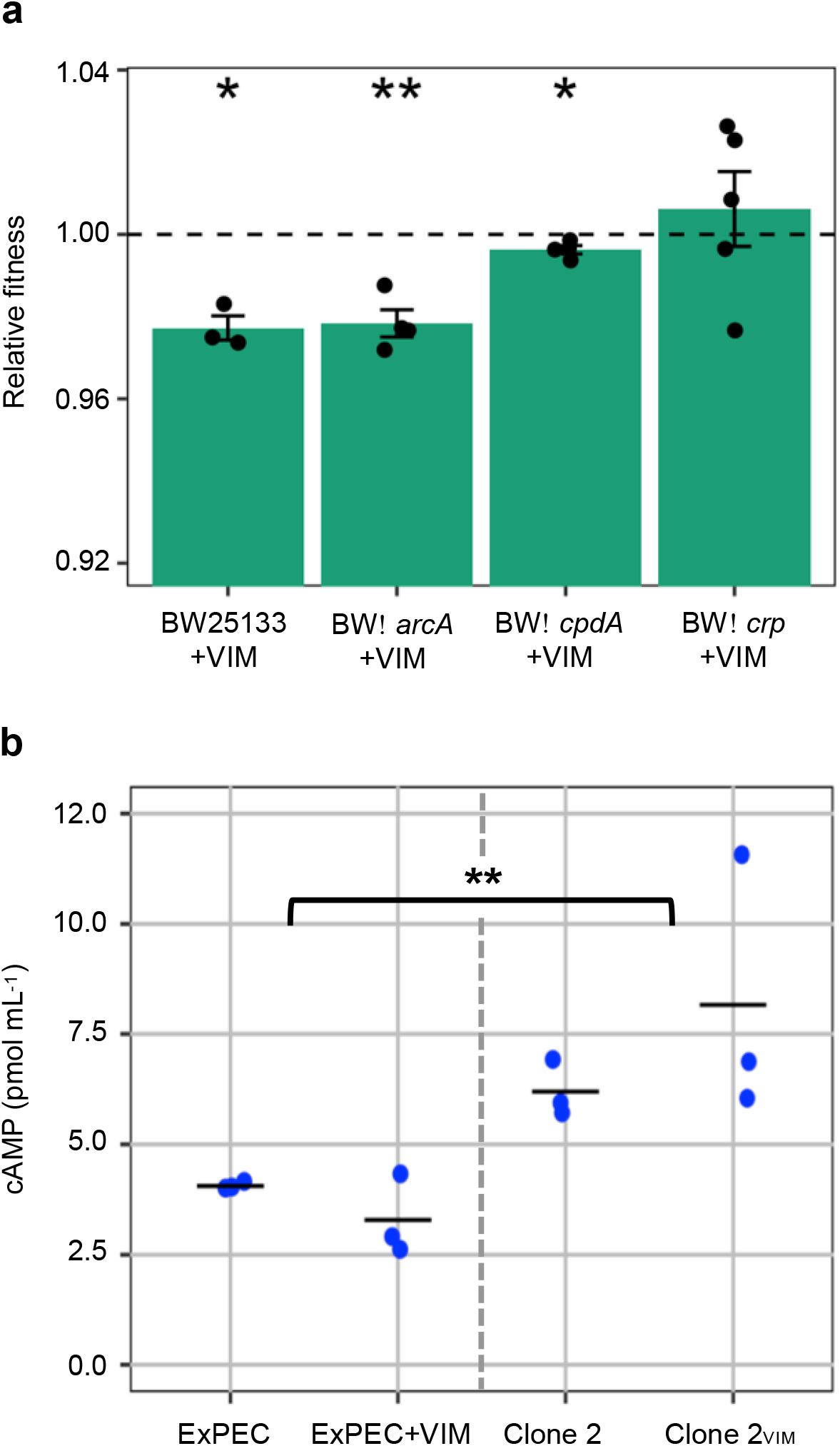
Effect of CCR and ArcAB systems mutations on plasmid cost and intracellular cAMP concentration. **a,** Fitness costs of pG06-VIM-1 in parent strain BW25133 and deletion strains (n = 3-5). Significant plasmid costs are indicated by asterisks (*P* = * < 0.05, ** < 0.01, *** < 0.001; one-sample *t*-test, two-sided). Error bars indicate ± s.e.m. **b** Intracellular cAMP concentrations of ancestral strains (ExPEC and ExPEC+VIM; n = 6; left) and evolved strains (Clone 2 and Clone 2_VIM_ carrying mutation *cpdA*.Δ3.bp488‑490; n = 6; right) (two-way ANOVA; *P* = ** < 0.01; df = 3).

In *E. coli*, the phosphodiesterase CpdA affects intracellular levels of cAMP by specifically hydrolyzing this signaling molecule [35]. To investigate the effect of the most frequently observed mutation in *cpdA* (*cpdA*.Δ3.bp488‑490; Supplementary Table 4) on protein function, we measured intracellular cAMP concentrations in ancestral and evolved strains with and without pG06-VIM-1. The levels of intracellular cAMP increased significantly by 49% between ancestral and evolved strains (ExPEC and ExPEC+VIM: 2.6 to 4.3 pmol mL^−1^, mean = 3.7 ± 0.72 pmol mL^−1^; Clone 2 and Clone 2_VIM_: 5.7 to 11.6 pmol mL^−1^, mean = 7.2 ± 2.2 pmol mL^−1^; two-way ANOVA with interactions: df = 3, *P* = 0.005 (assuming equal variances) and *P* = 0.001 (adjusted for unequal variances); Figure 4b), but were unaffected by plasmid presence (two-way ANOVA with interactions: df = 3, *P* = 0.53 (assuming equal variances) and *P* = 0.40 (adjusted for unequal variances)). These data are consistent with the previously observed CpdA deficiency of an identical *E. coli* mutant resulting in an approximate doubling of intracellular cAMP [37]. Similarly, protein function analysis of evolved population data indicates that the majority of mutations in *cpdA* lead to the loss of CpdA function (Supplementary information section VII and Supplementary Table 9).

### Mutations in CCR and ArcAB regulatory systems lead to general adaptation to the growth conditions

The gene products of *cyaA, cpdA*, *crp*, *arcA*, *arcB* can be associated with transcription in *E. coli* involving the global regulators CRP and ArcA [38]. cAMP is an important second messenger that binds to CRP [39] and the complex activates cAMP-dependent regulation of carbon source utilization via the CCR system [35, 36]. Intracellular levels of cAMP in *E. coli* are controlled by CyA (synthesis) and CpdA (degradation) [35]. Proteins ArcA and ArcB compose the ArcAB two-component regulatory system involved in respiratory and energy metabolism of *E. coli* [40, 41]. Mutations in CCR- and ArcAB-associated proteins may lead to growth optimization in varying environments due to adaptation in downstream transcriptional regulatory networks [31, 39, 42].

To verify that the mutations identified in the two regulatory systems increase fitness in the given *in vitro* environment, we assessed the fitness of pG06-VIM-1-containing and - free, evolved and ancestral strains by measuring exponential growth rates. Plasmid pG06-VIM-1 in Clones 1-4_VIM_ displayed no mutations after experimental evolution and this approach allowed us to directly measure the effects of the chromosomal mutations on general fitness. Growth rates of evolved strains were increased by 7-17% across all comparisons independent of presence or absence of the plasmid (one-sample *t*-test, one-sided: Clone 1: *w* = 1.11 ± 0.02; *P* = 0.022; Clone 2: *w* = 1.17 ± 0.03; *P* = 0.019; Clone 3: *w* = 1.13 ± 0.02; *P* = 0.009; Clone 4: *w* = 1.14 ± 0.02; *P* = 0.007; Clone 1_VIM_: *w* = 1.07 ± 0.02; *P* = 0.023; Clone 2_VIM_: *w* = 1.13 ± 0.06; *P* = 0.072; Clone 3_VIM_: *w* = 1.17 ± 0.05; *P* = 0.040; Clone 4_VIM_: *w* = 1.16 ± 0.04; *P* = 0.026; Figure 5a and b). Despite lower resolution than competition experiments, these data show that the identified mutations increase fitness under the given growth conditions and independent of plasmid carriage. They provide further support for the plasmid cost mitigating role of the observed chromosomal mutations.

**Figure 5.**
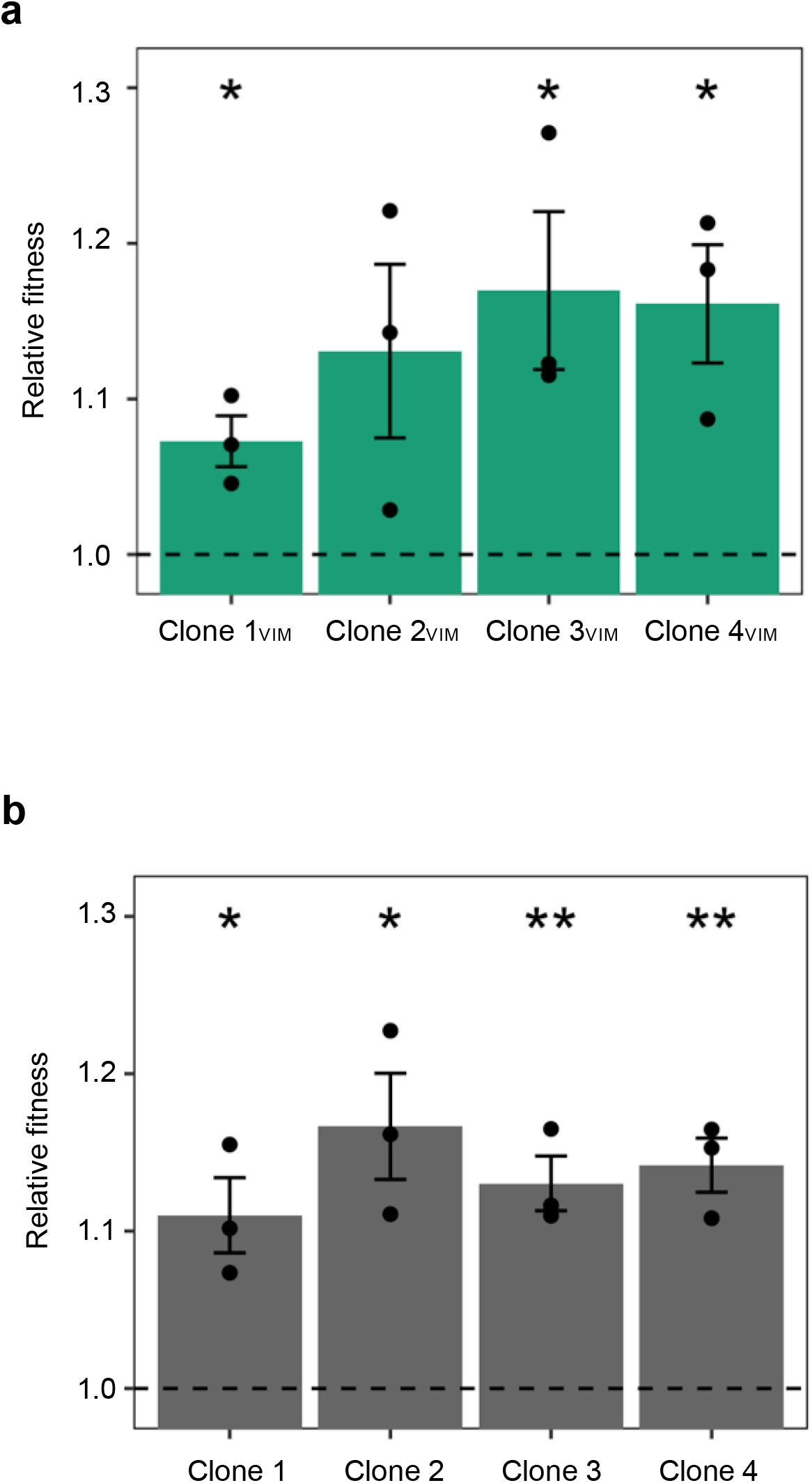
Fitness-improved adapted backgrounds. Relative fitness from comparison of exponential growth rates of **a** co-evolved pG06-VIM-1-carrying strains to strain ExPEC+VIM and (each comparison n = 3) **b** co-evolved pG06-VIM-1 segregants to ancestral strain ExPEC (each comparison n = 3). Significant fitness changes are indicated by asterisks (*P* = * < 0.05, ** < 0.01, *** < 0.001; one-sample *t*-test, one-sided). Error bars indicate ± s.e.m.

### Transcriptional alterations contribute to reduced plasmid costs

CRP and ArcA represent two of seven global transcription factors in *E. coli* and directly or indirectly control the expression of several hundred genes [43, 44]. Changes in gene expression may lead to a reduced burden of plasmid carriage, as demonstrated by Harrison *et al*. [20]. We sought to elucidate both the origin of the initial pG06-VIM-1 cost and its amelioration due to mutations in CCR and ArcAB associated genes, and performed RNA-Seq. Six replicate samples of plasmid-free strains ExPEC, Clone 2 and Clone 3, and plasmid-carrying strains ExPEC+VIM, Clone 2+VIM, Clone 3+VIM were sequenced resulting in on average 25 million paired-end reads per sample (Supplementary Table 10). Comparing the ancestral ExPEC strain with and without pG06-VIM-1 revealed that seven chromosomal genes were differently expressed upon plasmid acquisition, of which only the one encoding a putative selenium delivery protein displayed a fold-change beyond a 2-fold threshold (Figure 6a and Supplementary Table 11). The lack of substantial evidence for altered chromosomal gene expression suggests that the costly plasmid acquisition may be due to consumption of extra building blocks for plasmid replication and expression (e.g. ATP, nucleotides or amino acids [1]) instead of disruption of transcriptional regulation [45] or of specific cellular pathways e.g. SOS response [46].

**Fig. 6.**
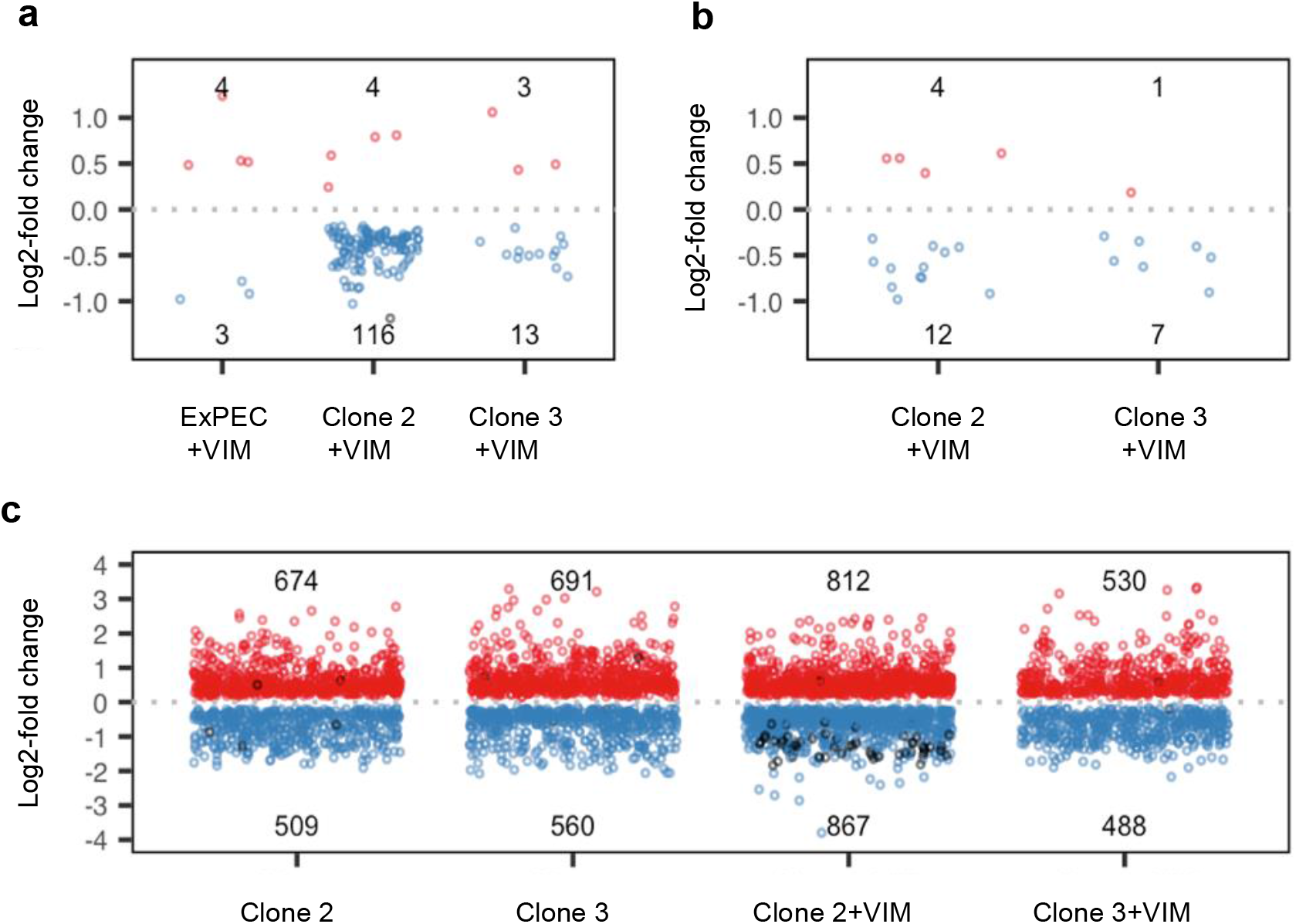
Differential expression analysis. Number of up and downregulated genes (Log_2_-fold change) on **a** chromosomes of ancestral strain ExPEC+VIM, evolved Clone 2+VIM and Clone 3+VIM upon acquisition of native pG06-VIM-1 (compared to respective plasmid-free strain) **b** evolved pG06-VIM-1 of Clone 2+VIM and 3+VIM due to adaptive chromosomal mutations (compared to ancestral ExPEC+VIM) **c** chromosomes of evolved pG06-VIM-1-free/-carrying Clone 2 and Clone 3 due to adaptive chromosomal mutations (compared to ExPEC and ExPEC+VIM, respectively). Circles: up (red) and downregulated (blue) protein-encoding genes; differently regulated RNA-encoding genes (black).

Given that the CCR and ArcAB systems are involved in global gene regulation it is not surprising that mutations in evolved Clone 2 and 3 lead to considerable changes in chromosomal gene expression when compared to the ancestral ExPEC strain. Indeed, hundreds of genes are differently up and downregulated independently of pG06-VIM-1 presence (Figure 6c; Supplementary Table 11). Despite some differences among the four evolved clones, enrichment (Supplementary Table 12) and over-representation (Supplementary Table 13) analyses of protein-encoding genes show common trends; cell motility via cilia/flagella (and other processes that require cell component organization, such as the expression of adhesion factors) tends to be downregulated (Supplementary Figure 5), while there is upregulation of diverse metabolic processes that target macromolecule biosynthesis (e.g. amino acids) and ribosome assembly which are, directly or indirectly, connected to translation and gene expression (Supplementary Figure 4).

After evolution, only 16 chromosomal genes of Clone 3 were affected by pG06-VIM-1 acquisition, of which one gene encoding a phage tail protein, exhibited overexpression > 2-fold (Figure 6a; Supplementary Table 11). In Clone 2 we found upregulation of four chromosomal genes while 116 were downregulated (Figure 6a; Supplementary Table 11). Analyzing the 115 downregulated protein-encoding genes revealed an over-representation of biological processes involved in tRNA metabolism and nucleotide biosynthesis (Supplementary Table 13; Supplementary Figure 6), while the remaining gene encodes a tRNA. Furthermore, the comparison Clone 2+VIM vs ExPEC+VIM revealed that 52 (or 65%) of the strain’s tRNA-encoding genes are downregulated (Figure 6c; Supplementary Table 11) which is indicative of altered translation processes. Therefore, at least in Clone 2 the low cost of pG06-VIM-1 can be attributable to interference in translation, which is in agreement with other reports showing that low plasmid costs are associated with gene expression [45, 47]. Interestingly, overall expression of pG06-VIM-1 genes decreased in evolved hosts, such that in Clone 2+VIM 12 plasmid genes are downregulated and four upregulated, while in Clone 3+VIM seven plasmid genes are downregulated but only one is upregulated (Figure 6b; Supplementary Table 11). Although transcriptional changes in these genes never exceed a 2-fold threshold, the net fold-change is negative (−13.08 for Clone 2+VIM and −9.01 for Clone 3+VIM; Supplementary Table 17). Taken together, these data suggest that net downregulation of plasmid genes after evolution offer a plausible explanation for the reduced fitness costs. Thus, reshaping gene expression at a global level through the identified mutations in the CCR and ArcAB systems affects plasmid transcription levels. This represents a novel solution to the plasmid paradox where adaptation to a new niche (growth medium in this case) pleiotropically mediates plasmid cost reductions.

## Discussion

In this report we asked if and how plasmid-host coevolution would mitigate the fitness costs of two clinically highly relevant MDR plasmids newly acquired by a plasmid-free ExPEC isolate. Our data show that the moderate initial costs of both carbapenemase-encoding plasmids were significantly alleviated during laboratory evolution. Curiously, these ameliorations of fitness costs were not due to plasmid-specific compensatory mutations as reported in several recent studies [10, 20, 21, 48] or deletions of costly plasmid regions in the larger conjugative plasmid [17]. Instead, after ~300 generations we identified strong parallel evolution in chromosomal genes only, independent of plasmid carriage. The mutational target genes represented two global regulatory systems involved in *E. coli* carbon catabolite repression (CCR) and aerobic respiration (ArcAB). Moreover, the mutations in these transcriptional regulators relieved the initial fitness costs of two unrelated plasmids strongly suggesting that the ExPEC host became generally more permissive towards plasmid acquisition. The pleiotropic effects on plasmid cost amelioration appears to be mainly due to mutations affecting the CCR regulatory system, as demonstrated by fitness results using *cpdA* and *crp* deletion mutants. Mechanistically, RNA-seq analyses revealed a net transcriptional relief on plasmid genes as a collateral cost-mitigating effect of environmental adaptation by global regulatory changes.

Other studies have also reported that mutations in regulatory systems improved plasmid-host relationships. In a seminal study, mutations in the *gacA/gacS* two-component regulatory system reduced the cost of the mega plasmid pQBR103 by decreasing plasmid transcriptional demand in *Pseudomonas fluorescens* [20]. These mutations were specifically ameliorating the cost of the plasmid since they did not appear in the plasmid-free evolved lineages (true compensatory mutations) [20]. This is categorically different from our findings since we observed that adaptation in the CCR and ArcAB regulatory systems was not specific to plasmid-carrying populations. Two other reports frequently identified mutations in regulatory systems across multiple plasmid-carrying evolving populations that improved plasmid maintenance [18, 48]. However, the absence of evolved plasmid-free lineages in these studies preclude direct comparisons with the results presented here.

On a broader perspective, our results warrant further research on plasmid-evolutionary dynamics in different *E. coli* lineages and sub-lineages to better understand why some of them appear to be more prone to acquire and maintain MDR plasmids [6, 47]. Available data support that plasmids of clinical origins rarely reduce fitness of clinical host strains [29, 30, 45, 49–51] to the extent often seen in the pioneering studies of plasmid-host compensatory evolution [10, 20, 22, 52]. It is also clear that this alone cannot explain how successful clone-plasmid associations emerge. From population genomic analyses McNally and co-workers demonstrated an association between mutations in regulatory regions of the high-risk *E. coli* ST131 subclade C and the accessory genome, including MDR plasmids. In their interpretation, this finding represented evidence of compensatory evolution towards MDR plasmid acquisition. However, taken together with a recent report showing that ExPEC ST131 has adapted to separate ecological niches at the sub-clade level, our results provide an alternative explanation [53]. Based on data presented here, it can be hypothesized that regulatory changes could also in part represent niche-adaptations that coincidentally facilitate MDR plasmid acquisition and maintenance.

Our study is not without limitations. We acknowledge that the pleiotropic effects on plasmid costs reported here may be specific to a single environment as others have reported that both fitness costs [32, 54] and compensatory evolution [55] are highly media-dependent. Consequently, the specific mutations reported here may be media-dependent, but the processes targeted (i.e. global gene regulation) are widely reported across different media, strains, and plasmids supporting the generality of our findings.

In this report, we propose “piggybacking” on niche-adaptation as a novel, not mutually exclusive, solution for the “plasmid paradox”. Our approaches also underscore the importance of using clinically relevant strains and plasmids to investigate the evolutionary dynamics of plasmid-mediated antibiotic resistance. This knowledge can be used jointly with data from molecular epidemiology to better predict future emergence of successful combinations of clones, sub-lineages and antibiotic resistance determinants.

## Methods

### Bacterial hosts, plasmids and culture conditions

Strains, plasmids and primers used in this study are listed in Supplementary Tables 1 and 2. The ancestral plasmid-free strain ExPEC was collected in Greece as part of the ECO-SENS study and represents an *E. coli* urinary tract infection isolate of ST537 tested by multi-locus sequence typing and phenotypically susceptible to 24 antibiotics tested by disc diffusion [28, 56] (Supplementary Table 1). Plasmids pG06-VIM-1 and pK71-77-1-NDM originated from a *K. pneumoniae* wound infection isolate [26] and an uropathogenic *E. coli* [27] and were introduced into ExPEC by electroporation or conjugation, respectively. Strains were grown at 37°C under aeration in Miller Difco Luria-Bertani liquid broth (LB; Becton, Dickinson and Co.) or on LB agar (LBA) containing additional Select agar (15 g L^−1^, Sigma-Aldrich). For selection of plasmid-carrying strains, media were supplemented with ampicillin (100 mg L^−1^; Sigma-Aldrich). See Supplementary information section I for more details on strains constructed in this study.

### Experimental evolution

Single colonies of strains ExPEC, ExPEC+VIM and ExPEC+NDM were used to initiate four independent lineages each. The 12 lineages were evolved in 1 mL of antibiotic-free LB medium using 2 mL-deep-96-well plates in checkered pattern (VWR International) and incubated at 37°C with 700 rpm constant shaking (Microplate Shaker TiMix 5, Edmund Bühler). In total, 48 transfers with estimated 6.6 generations between two transfers (~300 generations) were performed involving a 1:100 dilution of stationary-phase cultures into fresh LB every 12 hours (~10^7^ cells transferred). Endpoint populations (Pop 1-4_VIM_, Pop 5-8_NDM_ and Pop 9-12) and one representative clone per plasmid-carrying evolved population (Clones 1-4_VIM_ and Clones 5-8_NDM_) were stored at −80°C (Supplementary information section I and Supplementary Table I).

### Whole-genome sequencing

See Supplementary information section II for details on long-read sequencing and assembly of a closed reference genome of strain ExPEC (GenBank accession CP053079). For Illumina whole-genome sequencing, genomic DNA of ancestral strains ExPEC, ExPEC+VIM, ExPEC+NDM, eight evolved clones (Clones 1-4_VIM_, Clones 5-8_NDM_) and 12 evolved mixed populations (Pop 1-4_VIM_, Pop 5-8_NDM_, Pop 9-12) (Figure 1) was isolated using the GenElute Bacterial Genomic DNA Kit (Sigma-Aldrich). DNA-purity and - quantity was assessed using a NanoDrop ND-1000 spectrophotometer (Thermo Scientific). Short-read sequencing library preparation and sequencing was performed following manufacturers’ instructions at the Genomic Support Centre Tromsø, UiT The Arctic University of Norway. The Nextera XT DNA Library preparation kit (Illumina) was used with an input of 1 ng genomic DNA and dual indexes. Samples were sequenced on a NextSeq 550 instrument (Illumina) with 300 cycles (2 × 150 bp paired-end reads), and a NextSeq 500/550 mid-output flow cell was used for clonal samples. One entire high-output flow cell was explicitly used for the population samples aiming at deep coverage. We ran Trim Galore v0.5.0 with default settings to remove adapter sequences (CTGTCTCTTATA) and low-quality bases, and SPAdes v3.13.0 with read error correction [57, 58]. Trimmed and error corrected short-reads were controlled for adapters and quality score using FastQC v0.11.4 [59]. The raw sequence reads (long and short) of 24 libraries are available from the NCBI Sequence Read Archive (SRA, BioProject accession PRJNA630076).

### Short-read sequence analysis

We used the breseq computational pipeline v0.33.0 and v0.35.0 for prediction of mutations from clonal and population short-read sequencing data [60]. Preprocessed reads (see above) of all evolved populations were mapped against the reference genome of strain ExPEC (GenBank accession CP053079), and against plasmid sequences of pG06-VIM-1 (Pop 1-4_VIM_; GenBank accession KU665641 [29]) or pK71-77-1-NDM (Pop 5-8 _NDM_; GenBank accession CP040884 [30]) when appropriate. Breseq was run with default settings except for specifications when analyzing clonal sequencing data (‘consensus-mode’; ‘frequency-cutoff 0.9’; ‘minimum-variant-coverage 10’; ‘consensus-minimum-total-coverage 10’) or population sequencing data (‘polymorphism-mode’; ‘frequency-cutoff 0.01’; ‘minimum-variant-coverage 10’; ‘minimum-total-coverage 100’; ‘base quality score 20’). We focused on the identification of *de novo* single nucleotide substitutions, deletions, insertions and small indels by manually evaluating the predicted mutations from the breseq outputs (Supplementary information section IIIb). The use of short-read sequencing data bears an inherent limitation regarding the interpretation of chromosomal inversions, rearrangements and mutations in repeat regions due to misaligned reads, and these mutations were thus omitted from further analysis. Repeats were confirmed using ‘tandem repeat finder’ v4.09 [61] or by manually searching the reference genome for multiple alignment options. For population sequencing analysis, we report genetic changes as low as 1% mutation frequency considering that the mutated locus reached ≥ 10% mutation frequency at least in one of the evolved populations. Artemis v16.0.0 (http://sanger-pathogens.github.io/Artemis/), Gene Construction Kit v4.0.3 (Textco Biosoftware Inc.) and the Integrative Genomics Viewer v2.6.0 (http://software.broadinstitute.org/software/igv/) were used to support manual inspection of sequencing data.

### Competitive fitness and plasmid stability

The relative competitive fitness (*w*) of plasmid-carrying clones was determined in pairwise serial competition experiments (~40 generations) with the isogenic plasmid-free strain, as described before [24], with minor modifications. Briefly, pre-adapted cultures of each competitor were adjusted to the same OD_600_, mixed in a 1:1 ratio, and used to initiate 1 mL batch cultures at a density of ~10^7^ CFU (= T_0_), in antibiotic-free LB and 2 mL-deep-96-well plates in checkered pattern (VWR International). Plates were incubated at 37°C with 700 rpm shaking (Microplate Shaker TiMix 5, Edmund Bühler), and the cultures were diluted 1:100 into fresh LB every 12 hours (= T_12-72_). To determine the CFU of each competitor, cultures were diluted in 0.9% saline (m/v) and plated selectively on LBA-ampicillin (CFU_plasmid-carrying_) and non-selectively on LBA (CFU_total_) at T_0_ and every following timepoint. The selection coefficient was calculated as *s* = 0.5 × *b*/ln(1/*d*) with *b* (= slope) obtained from regressing the natural logarithm of the ratio (CFU_plasmid-carrying_/CFU_plasmid-free_) over timepoints, and *d* as the dilution factor at each transfer (here 1:100) [62]. It was multiplied by 0.5 to account for two transfers per day (to obtain *s* per day). Relative fitness was calculated as *w* = 1+*s*, where the fitness of the plasmid-free strain equals 1 (Supplementary Table 6). To determine spontaneous plasmid loss during competition experiments we proceeded similarly with pre-adapted cultures of plasmid-carrying strains as described above. Briefly, the density at T_0_ was ~5 × 10^6^ CFU mL^−1^ and cultures were transferred, diluted and plated selectively and non-selectively, as described above. The slope obtained by regressing the frequency of the plasmid-carrying population (CFU_plasmid-carrying_/CFU_total_) over timepoints was calculated (Supplementary Table 7). For determination of relative competitive fitness and spontaneous plasmid loss, results were obtained from at least three biological replicates, initiated on separate days, with three technical replicates each.

### Exponential growth rates

As a proxy for fitness changes due to acquired mutations in evolved clones with and without the plasmid, the exponential growth rates of separately growing strains were determined. Briefly, overnight cultures in 1 mL LB were started from a single colony grown on LBA, diluted 1:100 in LB, and 250 μl were aliquoted into a 96-well-microtiter plate (Thermo Scientific). Absorbance at OD_600_ nm was measured in a BioTek EPOCH2 microtiter spectrophotometer (BioTek Instruments), every 10 minutes, and with linear shaking. Growth rates (*r*) were determined using GrowthRates v3.0 [63]. Fitness of the evolved strain was calculated as relative growth rate = *r*_evolved strain_/*r*_ancestral strain_. Results were obtained from three biological replicates including five technical replicates all displaying a correlation coefficient R ≥ 0.97.

### Intracellular cyclic adenosine monophosphate (cAMP) concentration

Intracellular cAMP was quantified using the Cyclic AMP Select ELISA Kit (Cayman Chemical) following manufacturers’ instructions. For this purpose, overnight cultures were started from single colonies into 2 mL LB, diluted 1:100 into fresh LB and incubated until mid-exponential growth phase (between 5.3-6.7 × 10^8^ CFU mL^−1^). Five mL of each culture were spun down at 4°C, 4000 rpm for 10 min (Eppendorf Centrifuge 5810), supernatant was removed, and the pellet was subsequently washed three times in ice-cold 0.9% saline (m/v). Cells were resuspended in 250 μl of 0.1 M HCl to stop endogenous phosphodiesterase activity and the suspensions were boiled for 5 minutes. After centrifugation at 4°C, 4000 rpm for 10 min (Eppendorf Centrifuge 5810) an aliquot of the supernatant was diluted 1:2 (ExPEC, ExPEC+VIM) or 1:8 (Clone 2_VIM_, Clone 2) in ELISA buffer followed by a subsequent 1:2 dilution step for all samples. The cAMP standard was reconstituted in 0.1 M HCl but thereafter diluted into ELISA buffer. Samples were applied in two dilutions, each in three biological and three technical replicates on the same ELISA plate. The standard was applied once and in two technical replicates on the same plate. The plate was incubated in the dark for 18 hours at 4°C and thereafter developed under slow orbital shaking and dark conditions. Absorbance was measured at 410 nm periodically in a BioTek EPOCH2 microtiter spectrophotometer (BioTek Instruments). Final reads were taken at B_0-average_ = 0.7 and data was analyzed using the spreadsheet available at https://www.caymanchem.com/analysisTools/elisa (R^2^ standard curve = 0.96).

### Total RNA isolation

For transcriptome analysis, overnight cultures were initiated from single colonies into 2 mL LB, diluted 1:100 into fresh LB, and incubated until mid-exponential growth phase (OD_600_ 0.5-0.6; average 2.2±0.9 × 10^8^ CFU mL^−1^). Total RNA was isolated in six biological replicates per strain from 0.5 mL of culture using the RNeasy Protect Bacteria Mini kit (Qiagen) on six consecutive days. RNA-quality and −quantity were assessed with Nanodrop ND-1000 spectrophotometer (Thermo Scientific). Contaminating genomic DNA (gDNA) was digested following rigorous DNAse I treatment of the Ambion DNA-free DNase kit (Thermo Scientific). Briefly, 50 μl assays of maximum 10 μg RNA were treated in two consecutive incubation steps at 37°C for 30 minutes and addition of 5 μl DNase I enzyme before each step. RNA-quality and −quantity were again assessed as described above and the absence of gDNA was tested by PCR amplification (40 cycles) of the *adk* housekeeping gene (Supplementary Table 2). The RNA integrity numbers (RIN) were obtained via the Agilent RNA 6000 Nano kit and the Agilent 2100 Bioanalyzer system (Agilent Technologies 2100), and all samples reached RIN > 9 (Supplementary Table 10). Depletion of ribosomal RNA from 1 μg total RNA per sample with the QIAseq FastSelect RNA Removal kit and library preparation using the Truseq Stranded mRNA library kit were performed at Qiagen (Genomic Service Hilden, Germany). The Norwegian Sequencing Centre (NSC) (http://www.sequencing.uio.no) performed sequencing of the library on 1/2×SP Novaseq flow cell with 300 cycles (2 × 150 bp paired-end reads). The raw sequence reads of 36 libraries are available from NCBI SRA (BioProject accession PRJNA630076).

### RNA-Seq analysis

NSC performed initial filtering of raw reads including adapter trimming and removal of low-quality reads using BBMap v34.56 (therein BBDuk) [64]. NSC mapped clean and adapter removed reads against the merged version of the ExPEC chromosome and the pG06-VIM-1 sequence using Hisat2 v2.1.0 [65] and generated count tables using FeatureCounts v1.4.6-p1 [66], resulting in an average sample alignment of 65%; Supplementary Table 10). Count tables were used as input for the Differential Expression analysis (data normalization and statistical tests) performed in R version 4.0.2 [67] using the default script for SARTools version 1.7.3 [68] with default settings and strain ExPEC as reference.

PANTHER Generic Mappings of chromosomal genes were generated using the PANTHER HMM Scoring tool with the PANTHER HMM library Version 15.0 [69–72] (Supplementary Table 16) and functional classification of the PANTHER accessions was retrieved from the website (Supplementary Table 15). Tabular lists containing the gene ID, PANTHER accession and fold change for all differentially expressed chromosomal genes were uploaded as PANTHER Generic Mappings to http://pantherdb.org/ and ran for enrichment of PANTHER GO-Slim Biological Processes with FDR (false discovery rate) correction. Enrichment analysis (Supplementary Table 12) was performed for each of the comparisons in Supplementary Table 11, except ExPEC+VIM vs ExPEC. For the over-representation analyses, subsets of the same lists (up and downregulated genes only) were uploaded separately to http://pantherdb.org/. Over-representation analyses (Supplementary Table 13) was performed against the PANTHER Generic Mapping of all chromosomal protein-encoding genes (Supplementary Table 14), with Fisher exact test and FDR correction. All GO terms displaying significant enrichment or over-representation were compiled in three subsets (up and downregulated processes independent of plasmid presence, and downregulated processes due to plasmid presence) and used to generate Supplementary Figures 4-6, respectively, in Visualize [73] available at http://amigo.geneontology.org/visualize?mode=client_amigo.

### Statistical analyses

Statistical analyses were performed in R version 4.0.2 [67]. Samples were verified for normality with Shapiro-Wilk test and/or graphical visualization. Homogeneity of variances was tested with Levene’s test (from package car [74]) and/or graphical visualization. One-sample or two-sample comparisons were performed with Student t-tests. Packages sandwich [75], car [74] and multcomp [76] were required for ANOVA and multiple comparisons, respectively. Graphs in Figures 1, 3, 4, 5 and 6 were produced with packages ggplot2 [77], patchwork [78], ggthemes [79], and RColorBrewer [80]. Significance levels are indicated as: *P*-value * < 0.05; ** < 0.01; *** < 0.001. Packages openxlsx [81] and writexl [82] were used to read/write xlsx files. Packages data.table [83] and jsonlite [84] were required to generate Supplementary Tables 11-16.

## Acknowledgements

We thank Francois Pierre Alexandre Cléon for technical assistance, and Prof. Ruth H. Paulssen and Hagar Taman at UiT Genomics Support Centre Tromsø for valuable discussions about sequencing approaches and service regarding Illumina sequencing. The PacBio and RNA sequencing services were provided by the Norwegian Sequencing Centre (www.sequencing.uio.no), a national technology platform hosted by the University of Oslo and supported by the “Functional Genomics” and “Infrastructure” programs of the Research Council of Norway and the South-Eastern Regional Health Authorities. We thank Teodora Ribarska and Arvind Sundaram at NSC for excellent correspondence and bioinformatic advice. This work was supported by the Northern Norway Regional Health Authority (Grant no. SFP1168-14), and a joint grant from Northern Norway Regional Health Authority and UiT The Arctic University of Norway (project A23270).

## Author contributions

P.J.J. and Ø.S. conceived the project; J.K., J.G. and P.J.J designed and J.K. and J.G. performed wet-lab experiments; J.H. performed reference genome assembly and provided bioinformatic support; J.K., J.G. and J.H. analyzed the data; all authors interpreted and discussed the data; J.K., J.G. and P.J.J. wrote the manuscript; all authors reviewed the manuscript and approved the final version.

## Data availability

The genome of reference strain ExPEC is available at GenBank (accession CP053079) and raw DNA- and RNA-sequencing data are accessible at NCBI SRA (BioProject accession PRJNA630076). All other relevant data are available within this article, the Supplementary Information files, or from the corresponding author upon request.

## Supplemental file 1

**Supplementary Information - Extended Methods and Results**

I. **Strain construction:**

supplementary methods
II. **Long-read whole-genome sequencing and *de novo* reference assembly:**

supplementary methods and results
III. **Short-read whole-genome sequencing:**

supplementary methods

a. **Populations:** supplementary results
b. **Clones:** supplementary results
IV. **Competitive fitness and plasmid loss:**

supplementary results
V. **Measurement of pK71-77-1-NDM transfer frequency:**

supplementary methods and results
VI. **Antimicrobial susceptibility testing:**

supplementary methods and results
VII. **Protein function:**

supplementary methods and results

**Supplemental file 1: Supplementary Information - Tables and Figures**

Supplementary Table 1. Plasmids and strains used in this study.

Supplementary Table 2. Primers used in this study.

Supplementary Table 3. Annotated genes present in deleted regions of evolved pK71-77-1-NDM.

Supplementary Table 4. Mutations identified in evolved populations.

Supplementary Table 5. Descriptive statistics from clonal and population short-read whole-genome sequencing data alignment to sequences of ancestral reference strain ExPEC and the respective plasmid.

Supplementary Table 6. Summary of the results from linear regression analysis (serial competition experiments).

Supplementary Table 7. Summary of the results from linear regression analysis (plasmid loss).

Supplementary Table 8. Results from susceptibility testing of ancestral and evolved pK71-77-1-NDM-carrying clones by disc diffusion test.

Supplementary Table 9. Effect of non-synonymous mutations on protein function in evolved populations.

**Supplementary Tables 10-17 (Supplemental File 2; Excel sheet).** Results RNA-Seq. RNA-Seq samples: quality and libraries statistics (Supp.Table 10); Differential Expression analysis (chromosome and plasmid) (Supp.Table 11); Enrichment analysis (chromosome) (Supp.Table 12); Over-representation analyses (chromosome) (Supp.Table 13); protein-encoding genes (chromosome) (Supp.Table 14); Functional Classification (chromosome) (Supp.Table 15); PANTHER Generic Mappings (Supp.Table 16); Net fold-change plasmid genes in Clone 2+VIM and Clone 3+VIM (Supp.Table 17).

Supplementary Figure 1. Read coverage plots of evolved pK71-77-1-NDM in evolved Pop 5_NDM_, Clone 5_NDM_ and Clone 7_NDM_.

Supplementary Figure 2. Number of chromosomal variants per evolved population and mutational target gene.

Supplementary Figure 3. Conjugation frequency of ancestral pK71-77-1-NDM from ancestral and evolved backgrounds into strain K56-43 RifR.

**Supplementary Figures 4-6 (additional PDF files).** Ontology graphs of enrichment and overrepresentation analysis RNA-Seq: upregulated processes in Clone 2 and 3 with and without pG06-VIM-1 due to adaptive mutations (Supp.Figure 4), downregulated processes in Clone 2 and 3 with and without pG06-VIM-1 due to adaptive mutations (Supp.Figure 5); downregulated processes due to plasmid presence in Clone 2+VIM (Supp.Figure 6).

