## Supplemental File 1 for "Piggybacking on niche-adaptation reduces the cost of multidrug resistance plasmids"

### Piggybacking on niche-adaptation reduces the cost of multidrug resistance plasmids

Julia Kloos[1#] (ORCID: 0000-0002-8773-3463), João A. Gama[1#] (ORCID: 0000-0003-0967-6616), Joachim Hegstad[1,2] (ORCID: 0000-0001-9173-7195), Ørjan Samuelsen[1,3] (ORCID: 0000-0002-5525-2614), Pål J. Johnsen[1*] (ORCID: 0000-0002-6455-6433)

[1]Department of Pharmacy, Faculty of Health Sciences, UiT The Arctic University of Norway, Tromsø, Norway; [2]Department of Microbiology and Infection Control, University Hospital of North Norway, Tromsø, Norway; [3]Norwegian National Advisory Unit on Detection of Antimicrobial Resistance, Department of Microbiology and Infection Control, University Hospital of North Norway, Tromsø, Norway.

[#]These authors contributed equally to this work.

[*]Corresponding author:
UiT The Arctic University of Norway
Department of Pharmacy
Faculty of Health Sciences
9037 Tromsø
Norway


**Supplementary Information - Extended Methods and Results**

I. **Strain construction:**

supplementary methods

II. **Long-read whole-genome sequencing and *de novo* reference assembly:**

supplementary methods and results

III. **Short-read whole-genome sequencing:**

supplementary methods

a. **Populations:** supplementary results

b. **Clones:** supplementary results

IV. **Competitive fitness and plasmid loss:**

supplementary results

V. **Measurement of pK71-77-1-NDM transfer frequency:**

supplementary methods and results

VI. **Antimicrobial susceptibility testing:**

supplementary methods and results

VII. **Protein function:**

supplementary methods and results

#### I. Strain construction.

See Supplementary Table 2 for all primers used during strain construction.

Supplementary methods:

Clones 1-4$_{\text{VIM}}$ and Clones 5-8$_{\text{NDM}}$: these strains are purified representatives of evolved, plasmid-carrying populations. Briefly, ampicillin-resistant colonies of increased size were screened for the presence of mutations in genes *arcA*, *arcB*, *cpdA* and *crp* by Sanger sequencing. For this purpose, colony PCR was performed using DreamTaq PCR master mix 2x (Thermo Scientific) with standard conditions. The PCR products were enzymatically purified using Exonuclease I (Thermo Scientific) and Shrimp Alkaline Phosphatase (rSAP; New England Biolabs). The purified amplicons were Sanger sequenced by BigDye 3.1 chemistry (Applied Biosystems) employing single primers of the respective primer-pair. The assays were analyzed in-house on an Applied Biosystems 3130*xl* Genetic Analyzer automated cycler.

Clones 1-4: these strains are spontaneous pG06-VIM-1-free segregants identified as ampicillin-susceptible isolates by screening co-evolved Clones 1-4$_{\text{VIM}}$ on LBA and LBA-ampicillin (Luria-Bertani agar plates without and with ampicillin). The presence of clone-specific mutations in Clones 1-4 was confirmed by Sanger sequencing as described above. However, no plasmid-free segregants of co-evolved Clones 5-8$_{\text{NDM}}$ were obtained using the same method.

Clone 2+VIM, Clone 3+VIM, BW25113+VIM, BWΔ*cpdA*+VIM, BWΔ*arcA*+VIM and BWΔ*crp*+VIM: these strains were either plasmid-free evolved segregants (Clone 2 and Clone 3) or deletion mutants from the Keio collection [1] that received the non-conjugative pG06-VIM-1 by electroporation. For this purpose, ancestral pG06-VIM-1 was isolated from ExPEC+VIM following the Qiagen protocol for isolation of very low-copy number plasmids using 500 mL culture volume and Qiagen-tip 100 (Qiagen). Electroporation was performed according to [2] and transformants were selected on LBA-ampicillin. Presence of pG06-VIM-1 was verified by replicon PCR (*repB*) in all purified clones. Sanger sequencing (see above) confirmed clone-specific mutations (Clone 2+VIM and Clone 3+VIM). PCR amplification and gel electrophoresis confirmed the deletion constructs of *cpdA, crp* and *arcA* in the Keio strains.

ExPEC+NDM, Clone 2+NDM and Clone 3+NDM: these strains carried the ancestral pK71-77-1-NDM and were obtained through conjugation by filter mating. For this purpose, M9 Minimal Medium agar was prepared from 5×M9 salts (200 mL $L^{-1}$) [3], 20% arabinose (20 mL $L^{-1}$), 200 mM $CaCl_2$ (0.5 mL $L^{-1}$), 1M $MgSO_4$ (2 mL $L^{-1}$), Select agar (15 g $L^{-1}$) and ampicillin (100 mg $L^{-1}$) (all Sigma-Aldrich). Tetrazolium Arabinose agar (TAA) was prepared from peptone (10 g $L^{-1}$; Oxoid, now Thermo Scientific), Yeast extract (0.1 g $L^{-}$1; Fluka, now Sigma-Aldrich), NaCl (5 g $L^{-1}$; Sigma-Aldrich), Select agar (15 g $L^{-1}$; Sigma-Aldrich), 20% arabinose (50 mL $L^{-1}$; Sigma-Aldrich) and 5% 2,3,5-Triphenyltetrazolium chloride (1 mL $L^{-1}$; Sigma-Aldrich). Each mating experiment was started by mixing overnight cultures of donor and recipient strains in a 1:1 ratio (500 µl each), 1:10 up-concentration of the mix by centrifugation, and application on a sterile filter (Pall Laboratory, mixed cellulose esters membrane) placed on LBA. After 6-8 hours of incubation at 37°C, bacteria were harvested by vortexing the filter in 2 mL of sterile 0.9% saline (m/v). Appropriate dilutions of the suspension were selectively plated. Firstly, strain K71-77 [4] served as the donor for ancestral pK71-77-1-NDM into rifampicin-resistant recipient strain MG1655 Δ*ara* RifR [5] generating strain MG1655 Δ*ara* RifR+NDM. Here, transconjugants carrying pK71-77-1-NDM were selected on LBA-rifampicin + ampicillin. A pure clone was verified for the inability to use arabinose as sole carbon source by a red colony phenotype on TAA. Presence of pK71-77-1-NDM ($bla_{NDM-1}$) and absence of pK71-77-2 (IncFIA-replicon) were confirmed by PCR. Secondly, MG1655 Δ*ara* RifR+NDM was employed as counter selectable pK71-77-1-NDM donor strain to generate transconjugants ExPEC+NDM, Clone 2+NDM and Clone 3+NDM, which were selected on M9 Minimal Medium agar containing arabinose and ampicillin. Pure clones were verified for their ability to use arabinose by a white colony phenotype on TAA, and by confirming the presence of pK71-77-1-NDM ($bla_{\text{NDM-1}}$) and clone-specific mutations by PCR and Sanger sequencing as described above.

#### II. Long-read whole-genome sequencing and *de novo* reference assembly.

Supplementary methods:

Long-read sequencing was performed on ancestral strain ExPEC (Supplementary Table 1) using Pacific Biosciences technology (Pacific Biosciences) at the Norwegian Sequencing Centre (http://www.sequencing.uio.no). For this purpose, genomic DNA was isolated from $4.5 \times 10^9$ cells (3.75 mL of a 4.0 McFarland suspension) of an overnight culture started from a single colony grown on LBA. Qiagen 100/G Genomic tips were used following the manufacturers' protocol for isolation of genomic DNA from Gram-negative bacteria (Qiagen). DNA-purity and -quantity were determined using NanoDrop ND-1000 spectrophotometer (Thermo Scientific). Additionally, DNA-quantity was assessed using Qubit High Sensitivity DNA assay (Thermo Scientific) and DNA-integrity was assessed on a 1% agarose gel. Long-read sequencing was performed using single-molecule real-time (SMRT) sequencing technology according to the Pacific Biosciences protocol with PacBio Barcoded Adapters for Multiplex Sequencing. The DNA was sheared to 20 kb fragments using Megaruptor (Diagenode). The final library was size selected using BluePippin (Sage Science) with 10 kb cutoff. The library was sequenced on a Pacific Biosciences Sequel instrument using Sequel Polymerase v3.0, SMRT cells v3 LR and Sequencing chemistry v3.0. Reads were demultiplexed using Barcoding pipeline on SMRT Link (v6.0.0.47841, SMRT

Link Analysis Services and GUI v6.0.0.47836) with 26 as minimum barcode score. Sequences were assembled using HGAP4 assembly pipeline on SMRT Link (v6.0.0.47841, SMRT Link Analysis Services and GUI v6.0.0.47836) with default settings and 5 MB as expected genome size.

We trimmed and downsized the long-read data to a total of 500 Mb using Filtlong v0.2.0 [6] whereby 10% of the worst-quality reads were discarded and the longest reads were maintained. A circularized reference genome of strain ExPEC was generated by *de novo* hybrid assembly of short- (see Supplementary information section IIIb) and long-read sequencing data using Unicycler v0.4.7 [7] with default settings. The circularized genome was corrected for single-nucleotide polymorphisms arising from alignment of the Illumina reads using Snippy v4.3.6 [8]. The final reference genome was annotated using the NCBI Prokaryotic Genome Annotation Pipeline [9] and is accessible at GenBank (accession CP053079).

#### III. Short-read whole-genome sequencing.

Supplementary methods:

Mutations from short-read sequencing data were predicted using the breseq computational pipeline v0.33.0 and v0.35.0 [10] as described in Methods. Further, breseq output files were manually screened; variants that were also predicted in the ancestral chromosome or plasmids when using the same alignment parameters (clonal or polymorphism mode) were subtracted from downstream analysis. Moreover, variants in homopolymeric regions or regions with ambiguous reference sequence due to poor sequencing coverage were excluded.

##### a) Populations.

Supplementary results:

We performed ‘deep’ short-read sequencing on all evolved populations (Pop 1-4$_{VIM}$, Pop 5-8$_{NDM}$, Pop 9-12). An average coverage of 1322×/1800× was reached for chromosomal or plasmid sequencing data, respectively (Supplementary Table 5). We did not observe signs of hypermutability from our population sequencing data and nonsense mutations were not reported for any of the evolved populations. In *cpdA* (3',5'-**c**yclic adenosine monophosphate (cAMP) **p**hospho**d**iesterase), genetic changes involved also small deletions ($\Delta$1, $\Delta$3, $\Delta$4 and $\Delta$13 bp) or short sequence insertions (+3, +4, +6 bp) (35%; Supplementary Table 4). Notably, mutations in *arcB* (**a**erobic **r**espiration **c**ontrol sensor protein; histidine kinase) were specific for one plasmid-free population (Pop 9) and three pK71-77-1-NDM-carrying populations (Pop 5$_{NDM}$, Pop 6$_{NDM}$ and Pop 7$_{NDM}$) (Supplementary Table 4). Details on variant position, type and frequency are listed in Supplementary Table 4.

The most abundant and usually also most frequent mutation present in CpdA was the deletion of three base pairs (TGC) in amino acid (aa) position 163 (*cpdA*.$\Delta$3.bp488-490) in 7/12 evolved populations. This deletion resulted in the loss of one leucine in a string of four leucines (aa 160-163). CpdA activity depends on the binding of two iron cations as cofactors and we identified aa substitutions and deletions in positions that are reported to be involved in metal cation binding in 4/12 evolved populations at low frequency (1.1-12.5%; D22A, H24Y/R, H164del (= *cpdA*.$\Delta$3.bp490-492), H203Y and H205Y) (UniProtKB-P0AEW4). Additionally, aa positions 24 and 205 of CpdA are described as binding sites for cAMP ligand to the enzyme and were affected by mutations in 3/12 evolved populations (UniProtKB-P0AEW4).

The predominant mutation in CRP (**c**AMP **r**eceptor **p**rotein; DNA-binding transcriptional regulator), G142S, was present in 5/10 evolved lineages, in which it was also the most frequent. The C-terminal part of CRP represents the DNA-binding HTH-domain (aa positions 138-210) and seven of in total eight identified mutations were located in this region of the protein (UniProtKB-P0ACJ8). In CRP, a hinge region located in aa positions 136-139 connects the N- and C-terminal domains of the protein [11], and aa changes in positions 142, 145, 149 and 135 in close proximity to this region are described to influence the cAMP dependence of the protein, especially when replaced with polar amino acids, resulting in cAMP-independent CRP* mutants [12] [13] [14]. Harman *et al.* described a CRP* phenotype of *E. coli* for aa substitution L196R [15], a variant which was also identified here. We did not identify mutations in aa positions that are, to our knowledge, directly involved in cAMP binding to CRP.

In ArcA (**a**erobic **r**espiration **c**ontrol protein; DNA-binding transcriptional regulator), mutation L50Q was the major variant and present in 5/12 populations but only most frequent in population Pop $3_{VIM}$ (100%). In this protein, a receiver domain is described for aa positions 1-135 and a DNA binding domain for aa positions 135-238 (UniProtKB-P0A9Q1). The majority of mutations identified in ArcA was located in the receiver domain (22/25), and only three in the DNA-binding domain, similar to mutations identified in *arcA* of *E. coli* and *Citrobacter freundii* adapting to nutrient-rich conditions [16].

The mutations identified in ArcB were specific to single evolved populations, except aa substitution S280R, which occurred in two populations. The majority of aa substitutions was identified in described regions of ArcB such as the PAS-domain (S159F), the PAC-domain (R269C), the primary transmitter domain (A410T, F413C, I442T), the receiver domain (L579W) and the secondary transmitter domain (L686F, F689V, L734R, C746W; UniProtKB-P0AEC3).

Additionally, single evolved populations obtained mutations in distinct chromosomal targets; a 15 bp-deletion in *polA* (DNA polymerase I; Pop 9) and a synonymous mutation in *yncE* (uncharacterized protein; Pop 4$_{\text{VIM}}$) (Supplementary Table 4). We found no indications that the mutation observed in Pop 2$_{\text{VIM}}$ upstream of *cyaA* affected the genes' promoter region.

None of the co-evolved populations had acquired point mutations in plasmid sequences. However, the plasmid coverage plot of evolved population Pop 5$_{\text{NDM}}$ displayed a considerable drop in read coverage depth for a ~8.8 kb region in pK71-77-1-NDM (Supplementary Figure 1) spanning from position ~77602 nt (*rmtC*; 16S rRNA-methyltransferase) to ~86464 nt (IS*1* family transposase) indicating a deletion. This plasmid segment contained an aminoglycoside resistance gene (*aac(3)-IId*), amongst other genes, but did not include the gene encoding the plasmids' carbapenemase (*bla*$_{\text{NDM-1}}$) (Supplementary Table 3).

**b) Clones.**

Supplementary results:

Short-read whole-genome sequencing of strains ExPEC, ExPEC+VIM, ExPEC+NDM, Clones 1-4$_{\text{VIM}}$ and Clones 5-8$_{\text{NDM}}$ resulted in an average coverage of 114×/130× for chromosomes and plasmids, respectively (Supplementary Table 5). Using short-read sequencing of strain ExPEC, MLST v2.0 [17], Plasmidfinder v2.0 [18] and Resfinder v4.0 [19] we confirmed that the reference strain belonged to sequence type 537 and was without plasmids or acquired antibiotic-resistance mutations. For all sequenced plasmid-carrying clones, pG06-VIM-1 and pK71-77-1-NDM copy-numbers were calculated by dividing plasmid coverage with chromosomal coverage obtained from short-read sequencing analysis (Supplementary Table 5). Plasmid copy-numbers were unchanged after evolution (Clones 1-4$_{\text{VIM}}$ = 0.9-1.5, average = 1.1; Clones 5-8$_{\text{NDM}}$ = 1.2-1.5, average = 1.4; while 1.3 for ExPEC+VIM and ExPEC+NDM). Read alignment of Clone 5$_{\text{NDM}}$ against ancestral pK71-77-1-NDM confirmed an ~8.8 kb deletion as described before for Pop 5$_{\text{NDM}}$ (Supplementary information section IIIa). Furthermore, an even larger plasmid deletion was observed from coverage plots for Clone $7_{NDM}$ covering a 58.9 kb region from position ~27596 nt (IS*1380*-like element; IS*Ec9* family transposase) to ~86456 nt (IS*1* family transposase) (Supplementary Figure 1). The missing plasmid segment in Clone $7_{NDM}$ included the region that was also deleted in Clone $5_{NDM}$ in addition to genes of the adjacent deleted region (Supplementary Table 3). The deleted plasmid segments were flanked by mobile elements either on both sides (Clone $7_{NDM}$; IS*1380* and IS*1*) or at one side (Clone $5_{NDM}$; IS*1*) which suggests that recombination events involving these genetic elements lead to the deletion event.

#### IV. Competitive fitness and plasmid loss.

Supplementary results:

A summary of the results from linear regression analysis in serial competition experiments and from determination of plasmid loss is given in Supplementary Tables 6 and 7.

#### V. Measurement of pK71-77-1-NDM transfer frequency.

Supplementary methods:

12-hours-conjugation assays were performed to estimate interference of pK71-77-1-NDM transfer during fitness measurements. Ancestral strain ExPEC+NDM and evolved Clone 2+NDM and Clone 3+NDM represented donors for mating cultures. Rifampicin-resistant strain K56-43 RifR [4] was employed as recipient (Supplementary Table 1). Overnight cultures in LB were started for the respective strains and used to inoculate 1 mL LB with ~$10^7$ CFU $mL^{-1}$ of donor and recipient cells (1:1) in a 2 mL-deep-96-well plate (VWR International) (= $T_0$). The plate was incubated at 37°C and 700 rpm shaking

(Microplate Shaker TiMix 5, Edmund Bühler) for 12 hours (= $T_{12}$). Rifampicin (100 mg $L^{-1}$; Sigma-Aldrich) was added to LBA when appropriate. At $T_0$ and $T_{12}$, $CFU_{donor}$ and $CFU_{recipient}$ were determined on LBA-ampicillin and LBA-rifampicin, respectively. At $T_{12}$, $CFU_{transconjugants}$ was additionally determined on LBA-rifampicin + ampicillin. The frequency (%) of transconjugants per donor was calculated as:

$$\frac{CFU_{transconjugants}}{CFU_{donor}} \times 100$$

Supplementary results:

Results were obtained from three biological replicates with three technical replicates each. The determined conjugation frequency for ancestral pK71-77-1-NDM was as low as 0.22% (ExPEC+NDM), 0.09% (Clone 2+NDM) and 0.14% (Clone 3+NDM) (one-way ANOVA, df = 2, *P* = 0.058, not assuming equal variances; Supplementary Figure 3).

#### VI. Antimicrobial susceptibility testing.

Coverage plots from clonal short-read sequencing analysis indicated that deletions had occurred in co-evolved pK71-77-1-NDM sequences of Clone $5_{NDM}$ and Clone $7_{NDM}$ (Supplementary information section IIIb and Supplementary Figure 1). For both clones, these deletions included aminoglycoside resistance genes (Supplementary Table 3). To phenotypically examine these predicted deletions, we compared changes in the susceptibility of the ancestral strain ExPEC+NDM and the evolved Clones $5\text{-}8_{NDM}$ for two aminoglycoside antibiotics, tobramycin and gentamicin by disc diffusion.

Supplementary methods:

Antimicrobial susceptibility of ancestral strain ExPEC+NDM and evolved Clones 5-$8_{NDM}$ was investigated by disc diffusion tests on Mueller Hinton II agar (MHA; Becton,

Dickinson and Company). *E. coli* ATCC 25922 was included for quality control purposes. Briefly, 0.5 McFarland solutions in 0.9% saline (m/v) were prepared from freshly grown colonies on LBA, and the inocula were spread evenly onto MHA plates. One disc was applied per plate and strain for each antibiotic (10 µg per disc; Oxoid, now Thermo Scientific). Plates were incubated at 37°C and zone diameter was measured after 18 hours. Disc tests were interpreted (sensitive or resistant) according to EUCAST zone diameter breakpoints [20] (Supplementary Table 8). Measurements were repeated three times per drug and clone and we used the median of the results as final value for interpretation.

Supplementary results:

We found that strain ExPEC+NDM, Clone $6_{NDM}$ and Clone $8_{NDM}$ were fully resistant to these aminoglycosides, while Clone $5_{NDM}$ and Clone $7_{NDM}$ displayed increased susceptibility for both antibiotics. Both, Clone $5_{NDM}$ and Clone $7_{NDM}$ were now susceptible to gentamicin whereas only Clone $7_{NDM}$, which was affected by the larger deletion in pK71-77-1-NDM, was also susceptible to tobramycin (Supplementary Table 8). The changed resistance levels of Clone $5_{NDM}$ and Clone $7_{NDM}$ confirmed that deletions of the described pK71-77-1-NDM-regions had occurred during experimental evolution, which were likely mediated by plasmid-encoded mobile elements as also observed by Porse *et al.* [21].

#### VII. Protein function.

Supplementary methods:

Three online tools (PROVEAN (http://provean.jcvi.org/index.php) [22], SIFT (https://sift.bii.a-star.edu.sg) [23] and SNAP2 (https://rostlab.org/services/snap2web/) [24] were used to predict the effect of non-synonymous mutations, identified from clonal and whole population sequencing, on CpdA, CRP, ArcA and ArcB protein function. To conclude on a potential effect of a mutation, two of three prediction tools had to meet the criteria for ‘deleterious’ (PROVEAN), ‘effect’(SNAP2) or ‘affect protein function’(SIFT).

Supplementary results:

A majority (76%) of the 59 examined mutations were predicted to have a deleterious effect on protein function, and each prediction was either supported by all three (67% or 30/45) or at least two prediction models (33% or 15/45, Supplementary Table 9). For single proteins, 93% of the mutations were predicted deleterious in CpdA, 83% in ArcB, 68% in ArcA, and 50% in CRP.

#### Supplementary Information - Tables and Figures

Supplementary Table 1. Plasmids and strains used in this study.

Supplementary Table 2. Primers used in this study.

Supplementary Table 3. Annotated genes present in deleted regions of evolved pK71-77-1-NDM.

Supplementary Table 4. Mutations identified in evolved populations.

Supplementary Table 5. Descriptive statistics from clonal and population short-read whole-genome sequencing data alignment to sequences of ancestral reference strain ExPEC and the respective plasmid.

Supplementary Table 6. Summary of the results from linear regression analysis (serial competition experiments).

Supplementary Table 7. Summary of the results from linear regression analysis (plasmid loss).

Supplementary Table 8. Results from susceptibility testing of ancestral and evolved pK71-77-1-NDM-carrying clones by disc diffusion test.

Supplementary Table 9. Effect of non-synonymous mutations on protein function in evolved populations.

Supplementary Tables 10-17 (Supplemental File 2; Excel sheet). Results RNA-Seq. RNA-Seq samples: quality and libraries statistics (Supp.Table 10); Differential Expression analysis (chromosome and plasmid) (Supp.Table 11); Enrichment analysis (chromosome) (Supp.Table 12); Over-representation analyses (chromosome) (Supp.Table 13); protein-encoding genes (chromosome) (Supp.Table 14); Functional Classification (chromosome) (Supp.Table 15); PANTHER Generic Mappings (Supp.Table 16); Net fold-change plasmid genes in Clone 2+VIM and Clone 3+VIM (Supp.Table 17).

Supplementary Figure 1. Read coverage plots of evolved pK71-77-1-NDM in evolved Pop $5_{NDM}$, Clone $5_{NDM}$ and Clone $7_{NDM}$.

Supplementary Figure 2. Number of chromosomal variants per evolved population and mutational target gene.

Supplementary Figure 3. Conjugation frequency of ancestral pK71-77-1-NDM from ancestral and evolved backgrounds into strain K56-43 RifR.

Supplementary Figures 4-6 (additional PDF files). Ontology graphs of enrichment and overrepresentation analysis RNA-Seq: upregulated processes in Clone 2 and 3 with and without pG06-VIM-1 due to adaptive mutations (Supp.Figure 4), downregulated processes in Clone 2 and 3 with and without pG06-VIM-1 due to adaptive mutations (Supp.Figure 5); downregulated processes due to plasmid presence in Clone 2+VIM (Supp.Figure 6).

**Supplementary Table 1.** Plasmids and strains used in this study.

| **Plasmids**[a] | **Comment** | **Source** |
|---|---|---|
| pG06-VIM-1 | IncR; 53 kB; non-conjugative; *bla*$_{VIM-1}$, *aadA1, aadA2, aacA7, aphA, mphA, mphR, mrx, sul1, dfrA1, dfrA12, qacEΔ1*; originates from a *K. pneumoniae* wound infection isolate | [25], [26], GenBank KU665641 |
| pK71-77-1-NDM | IncC; 145 kB; conjugative; *bla*$_{NDM-1}$, *bla*$_{CMY-6}$, *aac(6')-Ib, aac(3)-II, rmtC, sul1, ble*$_{MBL}$; originates from an uropathogenic *E. coli* isolate | [27], [4], GenBank CP040884 |

| **Strains/Populations**[a] | **Comment** | **Source** | **Identifier**[b] |
|---|---|---|---|
| **Ancestral** | | | |
| ExPEC | *E. coli* K56-43; plasmid-free | [28], GenBank CP053079 | MP04-43 |
| ExPEC+VIM | ExPEC transformed with ancestral pG06-VIM-1; *E. coli* G1-15 in [26] | [26] | MP05-29 |
| ExPEC+NDM | ExPEC transconjugant harbouring pK71-77-1-NDM | this study | MP16-39 |
| **Evolved** | | | |
| Pop 1$_{VIM}$ | co-evolved ExPEC+VIM lineage 1; mixed population | this study | MP16-01 |
| Pop 2$_{VIM}$ | co-evolved ExPEC+VIM lineage 2; mixed population | this study | MP16-02 |
| Pop 3$_{VIM}$ | co-evolved ExPEC+VIM lineage 3; mixed population | this study | MP16-03 |
| Pop 4$_{VIM}$ | co-evolved ExPEC+VIM lineage 4; mixed population | this study | MP16-04 |
| Pop 5$_{NDM}$ | co-evolved ExPEC+NDM lineage 1; mixed population | this study | MP16-05 |
| Pop 6$_{NDM}$ | co-evolved ExPEC+NDM lineage 2; mixed population | this study | MP16-06 |
| Pop 7$_{NDM}$ | co-evolved ExPEC+NDM lineage 3; mixed population | this study | MP16-07 |
| Pop 8$_{NDM}$ | co-evolved ExPEC+NDM lineage 4; mixed population | this study | MP16-08 |
| Pop 9 | evolved ExPEC lineage 1; mixed population | this study | MP16-09 |
| Pop 10 | evolved ExPEC lineage 2; mixed population | this study | MP16-10 |
| Pop 11 | evolved ExPEC lineage 3; mixed population | this study | MP16-11 |
| Pop 12 | evolved ExPEC lineage 4; mixed population | this study | MP16-12 |
| Clone 1$_{VIM}$ | selected clone of Pop 1$_{VIM}$; ArcA.M39V; CpdA.S222L | this study | MP16-13 |
| Clone 2$_{VIM}$ | selected clone of Pop 2$_{VIM}$; ArcA.L117Q; *cpdA.*Δ3.bp488-490 | this study | MP16-14 |
| Clone 3$_{VIM}$ | selected clone of Pop 3$_{VIM}$; ArcA.L50Q; Crp.G142S | this study | MP16-15 |
| Clone 4$_{VIM}$ | selected clone of Pop 4$_{VIM}$; ArcA.F79Y; CpdA.P221L; YncE.T178T | this study | MP16-16 |
| Clone 5$_{NDM}$ | selected clone of Pop 1$_{NDM}$; ArcA.L93F; *cpdA.*Δ3.bp488-490 | this study | MP16-17 |
| Clone 6$_{NDM}$ | selected clone of Pop 2$_{NDM}$; ArcA.M39I; *cpdA.*Δ3.bp488-490 | this study | MP16-18 |
| Clone 7$_{NDM}$ | selected clone of Pop 3$_{NDM}$; ArcA.M53I; Crp.G142S | this study | MP16-19 |
| Clone 8$_{NDM}$ | selected clone of Pop 4$_{NDM}$; ArcA.M53I; Crp.T414R | this study | MP16-20 |
| **Strains derived from co-evolved clones** | | | |
| Clone 1 | plasmid-free segregant of Clone 1$_{VIM}$ | this study | MP16-21 |
| Clone 2 | plasmid-free segregant of Clone 2$_{VIM}$ | this study | MP16-22 |
| Clone 3 | plasmid-free segregant of Clone 3$_{VIM}$ | this study | MP16-23 |
| Clone 4 | plasmid-free segregant of Clone 4$_{VIM}$ | this study | MP16-24 |
| Clone 2+VIM | Clone 2 transformed with ancestral pG06-VIM-1 | this study | MP16-25 |
| Clone 3+VIM | Clone 3 transformed with ancestral pG06-VIM-1 | this study | MP16-26 |

| | | | |
|---|---|---|---|
| Clone 2+NDM | Clone 2 transconjugant harbouring pK71-77-1-NDM | this study | MP16-27 |
| Clone 3+NDM | Clone 3 transconjugant harbouring pK71-77-1-NDM | this study | MP16-28 |
| **Deletion strains** | | | |
| BW25113 | *E. coli* K-12 BW25113; parent strain of the Keio collection | [29] | MP16-31 |
| BWΔ*cpdA* | *E. coli* K-12 BW25113, Δ*cpdA*729::kan; Keio collection JW3000-1 | [1] | MP16-32 |
| BWΔ*arcA* | *E. coli* K-12 BW25113, Δ*arcA*726::kan; Keio collection JW4364-1 | [1] | MP16-33 |
| BWΔ*crp* | *E. coli* K-12 BW25113, Δ*crp*-765::kan; Keio collection JW5702-2 | [1] | MP16-34 |
| BW25113+VIM | BW25113 transformed with ancestral pG06-VIM-1 | this study | MP16-35 |
| BWΔ*cpdA*+VIM | BW25113 transformed with ancestral pG06-VIM-1 | this study | MP16-36 |
| BWΔ*arcA*+VIM | BW25113 transformed with ancestral pG06-VIM-1 | this study | MP16-37 |
| BWΔ*crp*+VIM | BW25113 transformed with ancestral pG06-VIM-1 | this study | MP16-38 |
| **Strains used in conjugation experiments** | | | |
| K56-43 RifR | spontaneous rifampicin-resistant mutant of *E. coli* K56-43 | [4] | MP16-29 |
| MG1655 Δ*ara* RifR | *E. coli* MG1655; Δ*ara*; rifampicin-resistant | [5] | MP03-11 |
| MG1655 Δ*ara* RifR+NDM | *E. coli* MG1655 Δ*ara*; rifampicin-resistant transconjugant harbouring pK71-77-1-NDM; counterselectable pK71-77-1-NDM donor | this study | MP16-30 |
| K71-77 | *E. coli*; donor of ancestral pK71-77-1-NDM; contains also pK71-77-2 | [4] | MP08-04 |

[a] plasmid/strain/population name in main article
[b] strain identifier at NCBI Sequence Read Archive, BioProject accession PRJNA630076

**Supplementary Table 2.** Primers used in this study.

| Target | Direction | Sequence (5'-3') | Comment | Source or reference |
|---|---|---|---|---|
| *repB* | F | TCGCTTCATTCCTGCTTCAGC | IncR replicon; characterization of pG06-VIM-1-transformants | [30] |
| | R | GTGTGCTGTGGTTATGCCTCA | | |
| *bla*$_{\text{NDM-1}}$ | F | CAGCAAATGGAAACTGGCGACCAA | carbapenemase gene; characterization of pK71-77-1-NDM-transconjugants | in-house |
| | R | ACGGTGATATTGTCACTGGTGTGG | | |
| iterons and *repE* | F | CCATGCTGGTTCTAGAGAAGGTG | IncFIA replicon (pK71-77-2); characterization of pK71-77-1-NDM-transconjugants | [31] |
| | R | GTATATCCTTACTGGCTTCCGCAG | | |
| *adk* | F | ATTCTGCTTGGCGCTCCGGG | *E. coli* house-keeping gene; confirmation absence of genomic DNA during RNA isolation | [32] |
| | R | CCGTCAACTTTCGCGTATTT | | |
| *arcA* | F | TGGAAAGTGCATCAAGAACG | mutational target gene; verification of identified mutations; verification of Δ*arcA* | this study |
| | R | TTCACTGCCGAAAATGAAAG | | |
| *arcB* | F_1 | GCAGGTTGTCGTGAAGGAAT | mutational target gene; verification of identified mutations | this study |
| | R_1 | TTACCGCCATGACTGTCTTTC | | |
| | F_2 | GCTGTCACGTTGGGGAATA | | |
| | R_2 | GTCTAGCCGGGGTCATTTTT | | |
| *cpdA* | F | GAAGTGTGTTCAAGCCAGCA | mutational target gene; verification of identified mutations; verification of Δ*cpdA* | this study |
| | R_1 | CCAGTTTACGTTCCAGCCAC | | |
| | R_2 | TGACGAACCGACAATACCCA | | |
| *crp* | F | AGCATATTTCGGCAATCCAG | mutational target gene; verification of identified mutations; verification of Δ*crp* | this study |
| | R | TAAATCAGTCTGCGCCACAT | | |
| *yncE* | F | GCATTGTGAAGCAGCAAAAA | verification of identified mutation in Clone $4_{\text{VIM}}$ | this study |
| | R | CGCGTCAATCACACTGACTT | | |

**Supplementary Table 3.** Annotated genes present in deleted regions of evolved pK71-77-1-NDM.

| Position | Gene/locus tag | Product | Deleted in |
|---|---|---|---|
| 27576..28838 | FHY70_00185 | IS*1380*-like element IS*Ec9* family transposase | Clone $7_{NDM}$ |
| 29162..30307 | FHY70_00190 | class C beta-lactamase CMY-6 | Clone $7_{NDM}$ |
| 30401..30934 | FHY70_00195 | lipocalin family protein | Clone $7_{NDM}$ |
| 30931..31248 | FHY70_00200 | quaternary ammonium compound-resistance protein SugE | Clone $7_{NDM}$ |
| 31505..31930 | FHY70_00205 | LuxR family transcriptional regulator | Clone $7_{NDM}$ |
| 31976..37462 | FHY70_00210 | DUF4165 domain-containing protein | Clone $7_{NDM}$ |
| 37611..38318 | FHY70_00215 | DsbC family protein | Clone $7_{NDM}$ |
| 38315..40762 | *traC* | type IV secretion system protein TraC | Clone $7_{NDM}$ |
| 40777..41094 | FHY70_00225 | hypothetical protein | Clone $7_{NDM}$ |
| 41091..41621 | FHY70_00230 | S26 family signal peptidase | Clone $7_{NDM}$ |
| 41584..42849 | FHY70_00235 | conjugal transfer protein TraW | Clone $7_{NDM}$ |
| 42846..43517 | FHY70_00240 | EAL domain-containing protein | Clone $7_{NDM}$ |
| 43514..44521 | FHY70_00245 | conjugal transfer protein TraU | Clone $7_{NDM}$ |
| 44625..47432 | *traN* | conjugal transfer mating pair stabilization protein TraN | Clone $7_{NDM}$ |
| 47471..48331 | FHY70_00255 | hypothetical protein | Clone $7_{NDM}$ |
| 48454..49095 | FHY70_00260 | hypothetical protein | Clone $7_{NDM}$ |
| 49390..49710 | FHY70_00265 | hypothetical protein | Clone $7_{NDM}$ |
| 50014..50199 | FHY70_00270 | hypothetical protein | Clone $7_{NDM}$ |
| 50419..51387 | FHY70_00275 | CbbQ/NirQ/NorQ/GpvN family protein | Clone $7_{NDM}$ |
| 51398..52306 | FHY70_00280 | hypothetical protein | Clone $7_{NDM}$ |
| 52367..52897 | *ssb* | single-stranded DNA-binding protein | Clone $7_{NDM}$ |
| 52992..53981 | *bet* | phage recombination protein Bet | Clone $7_{NDM}$ |
| 54044..55054 | FHY70_00295 | endonuclease | Clone $7_{NDM}$ |
| 54960..55172 | FHY70_00300 | hypothetical protein | Clone $7_{NDM}$ |
| 55254..56531 | FHY70_00305 | DUF3150 domain-containing protein | Clone $7_{NDM}$ |
| 56616..58412 | FHY70_00310 | VWA domain-containing protein | Clone $7_{NDM}$ |
| 58474..58809 | FHY70_00315 | hypothetical protein | Clone $7_{NDM}$ |
| 59074..59508 | FHY70_00320 | hypothetical protein | Clone $7_{NDM}$ |
| 59563..59766 | FHY70_00325 | hypothetical protein | Clone $7_{NDM}$ |
| 59838..60443 | FHY70_00330 | 5'-deoxynucleotidase | Clone $7_{NDM}$ |
| 60436..60705 | FHY70_00335 | hypothetical protein | Clone $7_{NDM}$ |
| 60719..60937 | FHY70_00340 | hypothetical protein | Clone $7_{NDM}$ |
| 61011..61568 | FHY70_00345 | pyruvate dehydrogenase | Clone $7_{NDM}$ |
| 61643..62494 | FHY70_00350 | NgrC | Clone $7_{NDM}$ |
| 62953..63339 | FHY70_00355 | hypothetical protein | Clone $7_{NDM}$ |
| 63517..65244 | FHY70_00360 | DNA primase | Clone $7_{NDM}$ |
| 65231..65509 | FHY70_00365 | hypothetical protein | Clone $7_{NDM}$ |
| 65582..65803 | FHY70_00370 | hypothetical protein | Clone $7_{NDM}$ |
| 65985..66989 | FHY70_00375 | IS*110*-like element IS*4321* family transposase | Clone $7_{NDM}$ |
| 67068..70040 | FHY70_00380 | Tn*3* family transposase | Clone $7_{NDM}$ |
| 70043..70600 | FHY70_00385 | recombinase family protein | Clone $7_{NDM}$ |
| 70638..70961 | FHY70_00390 | transposase | Clone $7_{NDM}$ |

| | | | |
|---|---|---|---|
| 70906..71919 | *intI1* | class 1 integron integrase IntI1 | Clone $7_{NDM}$ |
| 72126..72704 | *aac(6')-Ib* | AAC(6')-Ib family aminoglycoside 6'-N-acetyltransferase | Clone $7_{NDM}$ |
| 72873..73220 | FHY70_00405 | quaternary ammonium compound efflux SMR transporter QacE delta 1 | Clone $7_{NDM}$ |
| 73214..74053 | *sul1* | sulfonamide-resistant dihydropteroate synthase Sul1 | Clone $7_{NDM}$ |
| 74180..74422 | FHY70_00415 | hypothetical protein | Clone $7_{NDM}$ |
| 74506..74805 | FHY70_00420 | DUF2293 domain-containing protein | Clone $7_{NDM}$ |
| 74919..75089 | FHY70_00425 | CopG family transcriptional regulator | Clone $7_{NDM}$ |
| 75422..75973 | FHY70_00430 | DNA cytosine methyltransferase | Clone $7_{NDM}$ |
| 76016..76210 | FHY70_00435 | hypothetical protein | Clone $7_{NDM}$ |
| 76316..76969 | FHY70_00440 | endonuclease III | Clone $7_{NDM}$ |
| 77067..77912 | *rmtC* | RmtC family 16S rRNA (guanine(1405)-N(7))-methyltransferase | Pop 5 $_{NDM}$<br>Clone $5_{NDM}$<br>Clone $7_{NDM}$ |
| 78222..78919 | FHY70_00450 | IS*1* family transposase | Pop 5 $_{NDM}$<br>Clone $5_{NDM}$<br>Clone $7_{NDM}$ |
| 78925..79071 | FHY70_00455 | transposase | Pop 5 $_{NDM}$<br>Clone $5_{NDM}$<br>Clone $7_{NDM}$ |
| 79204..80064 | FHY70_00460 | aminoglycoside N-acetyltransferase AAC(3)-IId | Pop 5 $_{NDM}$<br>Clone $5_{NDM}$<br>Clone $7_{NDM}$ |
| 80077..80619 | FHY70_00465 | tunicamycin resistance protein | Pop 5 $_{NDM}$<br>Clone $5_{NDM}$<br>Clone $7_{NDM}$ |
| 81101..81292 | FHY70_00470 | hypothetical protein | Pop 5 $_{NDM}$<br>Clone $5_{NDM}$<br>Clone $7_{NDM}$ |
| 81298..81543 | FHY70_00475 | hypothetical protein | Pop 5 $_{NDM}$<br>Clone $5_{NDM}$<br>Clone $7_{NDM}$ |
| 81594..82730 | FHY70_00480 | DUF3883 domain-containing protein | Pop 5 $_{NDM}$<br>Clone $5_{NDM}$<br>Clone $7_{NDM}$ |
| 82845..84216 | FHY70_00485 | IS*1182*-like element IS*Cfr1* family transposase | Pop 5 $_{NDM}$<br>Clone $5_{NDM}$<br>Clone $7_{NDM}$ |
| 84816..85235 | FHY70_00490 | ATP-binding protein | Pop 5 $_{NDM}$<br>Clone $5_{NDM}$<br>Clone $7_{NDM}$ |
| 85238..85672 | FHY70_00495 | hypothetical protein | Pop 5 $_{NDM}$<br>Clone $5_{NDM}$<br>Clone $7_{NDM}$ |
| 85658..85780 | FHY70_00500 | metal-dependent hydrolase | Pop 5 $_{NDM}$<br>Clone $5_{NDM}$<br>Clone $7_{NDM}$ |
| 85726..85845 | FHY70_00505 | IS*6* family transposase | Pop 5 $_{NDM}$<br>Clone $5_{NDM}$<br>Clone $7_{NDM}$ |
| 85901..86598 | FHY70_00510 | IS*1* family transposase | Pop 5 $_{NDM}$<br>Clone $5_{NDM}$<br>Clone $7_{NDM}$ |

**Supplementary Table 4.** Mutations identified in evolved populations[a, b].

| Gene | Position | Mutation | Annotation | Pop 1$_{VIM}$ | Pop 2$_{VIM}$ | Pop 3$_{VIM}$ | Pop 4$_{VIM}$ | Pop 5$_{NDM}$ | Pop 6$_{NDM}$ | Pop 7$_{NDM}$ | Pop 8$_{NDM}$ | Pop 9 | Pop 10 | Pop 11 | Pop 12 |
|---|---|---|---|---|---|---|---|---|---|---|---|---|---|---|---|
| *arcA* | 4289006 | G→A | G36D (GGC→GAC) | | | | | | 2,2 | | | | | | |
| | 4289014 | A→G | M39V (ATG→GTG) ‡ | 15,2 | | | | | | | | | | | |
| | 4289016 | G→A | M39I (ATG→ATA) ‡ | | 1,2 | | | | 15,6 | | | 2,2 | | | |
| | 4289048 | T→A | L50Q (CTG→CAG) | | | 100 | 7,2 | | | 1,2 | | 1,2 | 1,7 | | |
| | 4289058 | G→A | M53I (ATG→ATA) | | | | | | | 38,9 | 33,5 | | | | |
| | 4289068 | C→A | L57M (CTG→ATG) | | | | | | | | | | | 2,6 | |
| | 4289079 | A→C | K60N (AAA→AAC) | | | | | | | | | | 93 | | |
| | 4289126 | C→A | A76E (GCG→GAG) | | | | | 4,5 | | | | | | | |
| | 4289134 | T→C | F79L (TTC→CTC) ‡ | | 31,6 | | | | | | | 33,2 | | | |
| | 4289135 | T→A | F79Y (TTC→TAC) ‡ | | | | 91,4 | | 2,6 | | | 17,1 | 1,1 | | |
| | 4289168 | T→C | I90T (ATT→ACT) | | | | | | | | | 8,1 | | | |
| | 4289173 | G→A | G92S (GGC→AGC) | | | | | | 9,2 | | | | | | |
| | 4289176 | C→T | L93F (CTC→TTC) | | | | | 94,5 | | | | | | | |
| | 4289179 | G→A | E94K (GAA→AAA) | 81 | | | | | | | | | | | |
| | 4289210 | C→A | P104Q (CCG→CAG) | | | | | | | | 1,5 | | 3,4 | | |
| | 4289212 | T→C | F105L (TTC→CTC) | | | | | | | | 19,4 | 17,5 | | | |
| | 4289234 | T→G | I112S (ATT→AGT) ‡ | | | | | | | | | | | 4,9 | |
| | 4289235 | T→G | I112M (ATT→ATG) ‡ | | | | | | | | 12,6 | | | | |
| | 4289243 | G→T | R115L (CGC→CTC) | | | | | | 3,8 | | 30,6 | | | | |
| | 4289249 | T→A | L117Q (CTG→CAG) | | 59,5 | | | | | | | | | 11,8 | |
| | 4289252 | T→A | L118Q (CTG→CAG) | | 4,3 | | | | 1,5 | | | | | | |
| | 4289258 | G→A | R120H (CGT→CAT) | | | | | | | | | | | 75,1 | 100 |
| | 4289357 | G→T | G153V (GGC→GTC) | | | | | | | | | | | 5,1 | |
| | 4289384 | C→T | P162L (CCG→CTG) | | | | | | 3 | | | | | | |
| | 4289512 | A→C | I205L (ATC→CTC) | | | | | | 2,2 | | | 2,6 | | | |
| *arcB* | 559547 | T→G | Y11D (TAT→GAT) | | | | | | 1,5 | | | | | | |
| | 559992 | C→T | S159F (TCC→TTC) | | | | | | | | | 5,3 | | | |
| | 560321 | C→T | R269C (CGT→TGT) | | | | | | | | 2,1 | | | | |
| | 560356 | C→A | S280R (AGC→AGA) | | | | | | | | 1,2 | 3,7 | | | |
| | 560744 | G→A | A410T (GCC→ACC) | | | | | | 2,2 | | | | | | |
| | 560754 | T→G | F413C (TTC→TGC) | | | | | | | | | 3,4 | | | |
| | 560841 | T→C | I442T (ATT→ACT) | | | | | | | 47,1 | | | | | |
| | 561252 | T→G | L579W (TTG→TGG) | | | | | | 27 | | | | | | |
| | 561574 | A→C | L686F (TTA→TTC) | | | | | | 22,3 | | | | | | |
| | 561581 | T→G | F689V (TTT→GTT) | | | | | | | 3,1 | | | | | |
| | 561717 | T→G | L734R (CTG→CGG) | | | | | | | 7,2 | | | | | |
| | 561754 | T→G | C746W (TGT→TGG) | | | | | | | | | 2,4 | | | |
| *cpdA* | 731329 | A→C | D22A (GAC→GCC) | | | | | | | | | | | | 4,1 |
| | 731334 | C→T | H24Y (CAC→TAC) ‡ | | | | | | | | | 3,5 | | | |
| | 731335 | A→G | H24R (CAC→CGC) ‡ | | | | | | 1,7 | | | | | | |
| | 731385 | A→C | S41R (AGC→CGC) | | | | | | | | | 3,3 | | | |
| | 731533 | Δ1 bp | coding (269/828 nt) | | | 8,5 | | | | | | | | | |
| | 731704 | Δ4 bp | coding (440-443/828 nt) | | | | | | | | | | | 1,1 | |
| | 731742 | +TGC | coding (478/828 nt) | 7,5 | 5,6 | | | | 15,3 | | | | | 8,9 | |
| | 731743 | Δ3 bp | coding (488-490/828 nt) | | 64 | | | 100 | 25,4 | 2,1 | 38,1 | 48 | 1,7 | | |
| | 731754 | Δ3 bp | coding (490-492/828 nt) | | | | | | | | | 2,3 | | | |
| | 731757 | C→T | H165Y (CAT→TAT) | | | | | | | 1,2 | | | | 1,6 | |
| | 731778 | Δ13 bp | coding (514-526/828 nt) | | | | | | | | 12,5 | | | | |
| | 731784 | T→G | W174G (TGG→GGG) | | | | | | 4,3 | | | | | | |
| | 731805 | +GTAA | coding (541/828 nt) | | | | 74,7 | | | | | | | | |
| | 731833 | T→A | L190Q (CTG→CAG) | | | | | | | | | | 1,8 | | |
| | 731863 | T→G | L200R (CTG→CGG) | | | | | | | | | 3,1 | | | |
| | 731868 | G→A | G202S (GGT→AGT) | | | | | | | | | | | | 70,2 |
| | 731869 | +TCACAT | coding (605/828 nt) | | | | | | 3,5 | | | | | | |
| | 731871 | C→T | H203Y (CAC→TAC) | | | | | | | | 12,5 | | | | |
| | 731877 | C→T | H205Y (CAT→TAT) | | | | | | | | 1,1 | | | | |
| | 731925 | C→A | P221T (CCG→ACG) ‡ | | | | | | | | 1,2 | | | | |
| | 731926 | C→T | P221L (CCG→CTG) ‡ | | | | 9,3 | | | | | | | | |
| | 731929 | C→T | S222L (TCG→TTG) | 13,6 | | | | | | | | | | | |
| | 732027 | A→C | T255P (ACC→CCC) | | | | | | | | | | | 32,9 | |
| *crp* | 427694 | A→C | L196R (CTG→CGG) | | | | | | | | | | 95,8 | | |
| | 427712 | A→C | M190R (ATG→AGG) | | | | | | | | | | | | 19,6 |
| | 427835 | A→C | L149R (CTG→CGG) | | | | | | | | | | | 4,1 | |
| | 427847 | G→A | A145V (GCA→GTA) ‡ | | | 9,9 | | | | | | | | 7,7 | |
| | 427848 | C→T | A145T (GCA→ACA) ‡ | | | | | | | | 4,2 | 34,1 | | | 1,5 |
| | 427857 | C→T | G142S (GGC→AGC) | 81,8 | | 77,3 | 7,5 | | 60,7 | 94,1 | | | | | |
| | 427859 | G→C | T141R (ACG→AGG) | | | | | | | | 33,2 | | | | |
| | 427878 | G→T | L135M (CTG→ATG) | | | | | | | | | | | 26,9 | |
| *cyaA ← / → hemC* | 5034928 | G→T | intergenic (-9/-378) | | 34,7 | | | | | | | | | | |
| *cyaA* | 5032557 | G→A | S788L (TCG→TTG) | | 1,6 | | | | | | | | | | |
| *polA* | 4951215 | +GCTGCACGTTCAGCC | coding (2529/2787 nt) | | | | | | | | | 19,9 | | | |
| *yncE* | 2595778 | G→A | T178T (ACC→ACT) | | | | 100 | | | | | | | | |

[a] ‡ = point mutations targeting the same codon
[b] grey background filling = mutations that were found in more than one evolved population

**Supplementary Table 5.** Descriptive statistics from clonal and population short-read whole-genome sequencing data alignment to sequences of ancestral reference strain ExPEC and the respective plasmid.

| Clone/Population | Read number | Mean coverage chromosome | Mean coverage plasmid[a] | Plasmid copy number[a] |
|---|---|---|---|---|
| ExPEC | 4196177 | 113 | n/a | n/a |
| ExPEC+VIM | 3242939 | 91 | 118 | 1,3 |
| ExPEC+NDM | 7187517 | 200 | 265 | 1,3 |
| Clone $1_{VIM}$ | 4778901 | 141 | 123 | 0,9 |
| Clone $2_{VIM}$ | 1715202 | 48 | 70 | 1,5 |
| Clone $3_{VIM}$ | 6000473 | 159 | 137 | 0,9 |
| Clone $4_{VIM}$ | 8490236 | 229 | 243 | 1,1 |
| Clone $5_{NDM}$ | 3731727 | 99 | 146 | 1,5 |
| Clone $6_{NDM}$ | 3451700 | 97 | 141 | 1,5 |
| Clone $7_{NDM}$ | 2486214 | 66 | 94 | 1,4 |
| Clone $8_{NDM}$ | 2539041 | 69 | 82 | 1,2 |
| **average** | **4149187** | **114×** | **130×** | |

| | | | |
|---|---|---|---|
| Pop $1_{VIM}$ | 24207523 | 650 | 1432 |
| Pop $2_{VIM}$ | 51031466 | 1412 | 2063 |
| Pop $3_{VIM}$ | 18387700 | 509 | 1132 |
| Pop $4_{VIM}$ | 22787328 | 644 | 363 |
| Pop $5_{NDM}$ | 57493948 | 1590 | 2428 |
| Pop $6_{NDM}$ | 37134452 | 1031 | 1446 |
| Pop $7_{NDM}$ | 58178599 | 1702 | 2217 |
| Pop $8_{NDM}$ | 78036459 | 2141 | 3322 |
| Pop 9 | 57282733 | 1611 | n/a |
| Pop 10 | 62460521 | 1894 | n/a |
| Pop 11 | 25260332 | 738 | n/a |
| Pop 12 | 63990083 | 1938 | n/a |
| **average** | **43407184** | **1322×** | **1800×** |

[a] n/a = not applicable

**Supplementary Table 6.** Summary of the results from linear regression analysis (serial competition experiments).

| Competitors | n | Mean relative fitness (*w*)[a] | SD[b] *w* | Shapiro–Wilk test | Min/max *w* | | Mean selection coefficient (*s*)[c] | *P s* | Significance *s*[d] | *P* Dunnet[e] | Significance Dunnet[e] |
|---|---|---|---|---|---|---|---|---|---|---|---|
| ExPEC vs ExPEC+VIM | 4 | 0.947 | 0.002 | 0.766 | 0.944 | 0.950 | -0.053 | $2.38 \times 10^{-5}$ | *** | control group | control group |
| Clone 1 vs Clone $1_{VIM}$ | 3 | 0.993 | 0.002 | 0.993 | 0.991 | 0.994 | -0.007 | 0.017 | * | $< 10^{-10}$ | *** |
| Clone 2 vs Clone $2_{VIM}$ | 3 | 0.996 | 0.001 | 0.336 | 0.995 | 0.997 | -0.004 | 0.016 | * | $< 10^{-10}$ | *** |
| Clone 3 vs Clone $3_{VIM}$ | 3 | 0.991 | 0.0004 | 0.122 | 0.990 | 0.991 | -0.009 | 0.001 | *** | $< 10^{-10}$ | *** |
| Clone 4 vs Clone $4_{VIM}$ | 3 | 0.994 | 0.002 | 0.271 | 0.992 | 0.996 | -0.006 | 0.056 | ns | $< 10^{-10}$ | *** |
| Clone 2 vs Clone 2+VIM | 3 | 0.987 | 0.004 | 0.290 | 0.985 | 0.992 | -0.013 | 0.026 | * | $2.51 \times 10^{-7}$ | *** |
| Clone 3 vs Clone 3+VIM | 3 | 0.990 | 0.001 | 0.365 | 0.989 | 0.990 | -0.010 | 0.002 | ** | $1.78 \times 10^{-7}$ | *** |
| ExPEC vs ExPEC+NDM | 3 | 0.945 | 0.012 | 0.627 | 0.932 | 0.956 | -0.055 | 0.017 | * | control group | control group |
| Clone 2 vs Clone 2+NDM | 3 | 0.976 | 0.007 | 0.553 | 0.971 | 0.984 | -0.024 | 0.025 | * | 0.006 | ** |
| Clone 3 vs Clone 3+NDM | 3 | 0.973 | 0.002 | 0.489 | 0.972 | 0.976 | -0.027 | 0.002 | ** | 0.010 | * |
| BW25113 vs BW25113+VIM | 3 | 0.977 | 0.005 | 0.238 | 0.974 | 0.983 | -0.023 | 0.016 | * | control group | control group |
| BWΔ*arcA* vs BWΔ*arcA*+VIM | 4 | 0.978 | 0.007 | 0.436 | 0.972 | 0.988 | -0.022 | 0.007 | ** | 0.993 | ns |
| BWΔ*cpdA* vs BWΔ*cpdA*+VIM | 4 | 0.996 | 0.002 | 0.750 | 0.994 | 0.999 | -0.004 | 0.034 | * | 0.0008 | *** |
| BWΔ*crp* vs BWΔ*crp*+VIM | 5 | 1.006 | 0.020 | 0.640 | 0.977 | 1.026 | 0.006 | 0.534 | ns | 0.051 | ns |

[a] relative fitness $w = 1+s$, where the fitness of the plasmid-free strain equals 1
[b] SD = standard deviation
[c] selection coefficient $s = 0.5 * b/\ln(1/d)$ with *b* (= slope) obtained from regressing the natural logarithm of the ratio ($CFU_{plasmid\text{-}carrying}/CFU_{plasmid\text{-}free}$) over timepoints (= $T_0$-$T_{72}$) and *d* as the dilution factor at each timepoint (here 1:100)
[d] one-sample *t*-test, two-sided; significance levels are indicated by asterisks (*P* = * < 0.05, ** < 0.01, *** < 0.001; ns = not significant)
[e] Dunnett's multiple comparison performed relative to the strain/strain-plasmid combination indicated as control group; significance levels are indicated by asterisks (*P* = * < 0.05, ** < 0.01, *** < 0.001; ns = not significant)

**Supplementary Table 7.** Summary of the results from linear regression analysis (plasmid loss).

| Strain | n | Mean slope (*b*)[a] | SD[*b*] *b* | *P b* | Significance *b*[c] | Mean frequency at $T_{72}$ | SD frequency at $T_{72}$ | *P* frequency at $T_{72}$ | Significance frequency[c] | Plasmid stability[d] |
|---|---|---|---|---|---|---|---|---|---|---|
| ExPEC+VIM | 3 | -0.011 | 0.002 | 0.017 | * | 0.967 | 0.065 | 0.477 | ns | stable |
| Clone $1_{VIM}$ | 3 | -0.012 | 0.026 | 0.501 | ns | 0.877 | 0.147 | 0.284 | ns | stable |
| Clone $2_{VIM}$ | 3 | 0.001 | 0.026 | 0.957 | ns | 0.849 | 0.052 | 0.037 | * | stable |
| Clone $3_{VIM}$ | 3 | -0.009 | 0.012 | 0.345 | ns | 0.790 | 0.048 | 0.017 | * | stable |
| Clone $4_{VIM}$ | 3 | -0.017 | 0.019 | 0.264 | ns | 0.892 | 0.116 | 0.248 | ns | stable |
| ExPEC+NDM | 3 | -0.006 | 0.011 | 0.458 | ns | 0.971 | 0.071 | 0.559 | ns | stable |
| Clone 2+VIM | 3 | -0.017 | 0.006 | 0.036 | * | 0.860 | 0.056 | 0.049 | * | unstable[e] |
| Clone 3+VIM | 3 | -0.004 | 0.027 | 0.824 | ns | 0.916 | 0.083 | 0.219 | ns | stable |
| Clone 2+NDM | 3 | 0.016 | 0.009 | 0.098 | ns | 0.998 | 0.027 | 0.893 | ns | stable |
| Clone 3+NDM | 3 | -0.009 | 0.016 | 0.459 | ns | 0.944 | 0.077 | 0.330 | ns | stable |
| BW25113+VIM | 3 | -0.015 | 0.059 | 0.702 | ns | 0.802 | 0.085 | 0.056 | ns | stable |
| BWΔarcA+VIM | 3 | 0.040 | 0.042 | 0.247 | ns | 0.998 | 0.143 | 0.981 | ns | stable |
| BWΔcpdA+VIM | 3 | 0.005 | 0.036 | 0.848 | ns | 1.103 | 0.113 | 0.254 | ns | stable |
| BWΔcrp+VIM | 3 | -0.012 | 0.042 | 0.665 | ns | 0.981 | 0.189 | 0.878 | ns | stable |

[a] slope obtained by regressing the frequency of the plasmid-carrying population ($CFU_{plasmid\text{-}carrying}$/ $CFU_{total}$) over timepoint (= $T_0$-$T_{72}$)
[b] SD = standard deviation
[c] one-sample *t*-test, two-sided; significance levels are indicated by asterisks (*P* = * < 0.05, ** < 0.01, *** < 0.001; ns = not significant)
[d] plasmids were considered stable if a *P*-value for either '*b*' or 'Mean frequency at $T_{72}$' was non-significant
[e] in Clone 2+VIM, 'Mean frequency at $T_{72}$' is only borderline significant

**Supplementary Table 8.** Results from susceptibility testing of ancestral and evolved pK71-77-1-NDM-carrying clones by disc diffusion test[a,b].

| | **Tobramycin (10 µg)** | **Gentamicin (10 µg)** |
|---|---|---|
| ATCC 25922[c] | 18 = S | 19 = S |
| | 19 = S | 19 = S |
| | 19 = S | 19 = S |
| | **19 = S** | **19 = S** |
| Clone $5_{NDM}$ | 8 = R | 17 = S |
| | 8 = R | 18 = S |
| | 7 = R | 18 = S |
| | **8 = R** | **18 = S** |
| Clone $6_{NDM}$ | no inhibition zone = R | no inhibition zone = R |
| | no inhibition zone = R | no inhibition zone = R |
| | no inhibition zone = R | no inhibition zone = R |
| | no inhibition zone = **R** | no inhibition zone = **R** |
| Clone $7_{NDM}$ | 20 = S | 20 = S |
| | 20 = S | 20 = S |
| | 20 = S | 21 = S |
| | **20 = S** | **20 = S** |
| Clone $8_{NDM}$ | no inhibition zone = R | no inhibition zone = R |
| | no inhibition zone = R | no inhibition zone = R |
| | no inhibition zone = R | no inhibition zone = R |
| | no inhibition zone = **R** | no inhibition zone = **R** |
| ExPEC+NDM | no inhibition zone = R | no inhibition zone = R |
| | no inhibition zone = R | no inhibition zone = R |
| | no inhibition zone = R | no inhibition zone = R |
| | no inhibition zone = **R** | no inhibition zone = **R** |

[a] S = susceptible; R = resistant (according to [20])
[b] median inhibition zone diameter (mm)
[c] control strain

**Supplementary Table 9.** Effect of non-synonymous mutations on protein function in evolved populations[a].

| | PROVEAN[b] | | SIFT[c] | | SNAP2[d] | | | overall effect[e] |
|---|---|---|---|---|---|---|---|---|
| | score | effect | score | effect | score | effect | expected accuracy | |
| ***arcA*** | | | | | | | | |
| **G36D** | -5.33 | Deleterious | 0.01 | AFFECT PROTEIN FUNCTION | 79 | effect | 85% | 3/3 |
| **M39V** | -2.95 | Deleterious | - | TOLERATED | 15 | effect | 59% | 2/3 |
| **M39I** | -2.916 | Deleterious | 0.32 | TOLERATED | 28 | effect | 65% | 2/3 |
| **L50Q** | -5.513 | Deleterious | 0 | AFFECT PROTEIN FUNCTION | 45 | effect | 71% | 3/3 |
| **M53I** | -0.87 | Neutral | 0.06 | TOLERATED | 32 | effect | 66% | 3/3 |
| **L57M** | -1.631 | Neutral | 0 | AFFECT PROTEIN FUNCTION | -93 | neutral | 97% | 2/3 |
| **K60N** | -2.679 | Deleterious | 0.14 | TOLERATED | 45 | effect | 71% | 2/3 |
| **A76E** | -2.102 | Neutral | 0.09 | TOLERATED | 70 | effect | 85% | 2/3 |
| **F79L** | 0.765 | Neutral | - | TOLERATED | -47 | neutral | 72% | 3/3 |
| **F79Y** | -1.549 | Neutral | 0.03 | AFFECT PROTEIN FUNCTION | 60 | effect | 80% | 2/3 |
| **I90T** | -4.928 | Deleterious | 0.01 | AFFECT PROTEIN FUNCTION | 50 | effect | 75% | 3/3 |
| **G92S** | -4.827 | Deleterious | 0 | AFFECT PROTEIN FUNCTION | 74 | effect | 85% | 3/3 |
| **L93F** | -3.93 | Deleterious | 0 | AFFECT PROTEIN FUNCTION | 1 | effect | 53% | 3/3 |
| **E94K** | -3.996 | Deleterious | 0 | AFFECT PROTEIN FUNCTION | 80 | effect | 91% | 3/3 |
| **P104Q** | -7.993 | Deleterious | 0 | AFFECT PROTEIN FUNCTION | 80 | effect | 91% | 3/3 |
| **F105L** | -4.943 | Deleterious | 0.41 | TOLERATED | 44 | effect | 71% | 2/3 |
| **I112M** | -1.414 | Neutral | 0.09 | TOLERATED | -63 | neutral | 82% | 3/3 |
| **I112S** | -2.444 | Neutral | 0.06 | TOLERATED | -26 | neutral | 61% | 3/3 |
| **R115L** | -5.594 | Deleterious | 0.01 | AFFECT PROTEIN FUNCTION | 61 | effect | 80% | 3/3 |
| **L117Q** | -5.665 | Deleterious | 0.01 | AFFECT PROTEIN FUNCTION | 13 | effect | 59% | 3/3 |
| **L118Q** | -4.794 | Deleterious | 0 | AFFECT PROTEIN FUNCTION | 21 | effect | 63% | 3/3 |
| **R120H** | -4.837 | Deleterious | 0 | AFFECT PROTEIN FUNCTION | 90 | effect | 95% | 3/3 |
| **G153V** | -1.529 | Neutral | 0.25 | TOLERATED | -22 | neutral | 61% | 3/3 |
| **P162L** | -3.961 | Deleterious | 0.06 | TOLERATED | 61 | effect | 80% | 2/3 |
| **I205L** | -1.88 | Neutral | 0 | AFFECT PROTEIN FUNCTION | -2 | neutral | 53% | 2/3 |
| ***arcB*** | | | | | | | | |
| **Y11D** | -6.971 | Deleterious | 0 | AFFECT PROTEIN FUNCTION | 79 | effect | 85% | 3/3 |
| **S159F** | -5.335 | Deleterious | 0.02 | AFFECT PROTEIN FUNCTION | 24 | effect | 63% | 3/3 |
| **R269C** | -6.825 | Deleterious | 0.01 | AFFECT PROTEIN FUNCTION | -18 | neutral | 57% | 2/3 |
| **S280R** | -4.062 | Deleterious | 0.04 | AFFECT PROTEIN FUNCTION | 21 | effect | 63% | 3/3 |
| **A410T** | -3.762 | Deleterious | 0.01 | AFFECT PROTEIN FUNCTION | 49 | effect | 71% | 3/3 |
| **F413C** | -7.599 | Deleterious | 0 | AFFECT PROTEIN FUNCTION | 20 | effect | 63% | 3/3 |
| **I442T** | -4.779 | Deleterious | 0.03 | AFFECT PROTEIN FUNCTION | -4 | neutral | 53% | 2/3 |
| **L579W** | -5.787 | Deleterious | 0 | AFFECT PROTEIN FUNCTION | 59 | effect | 75% | 3/3 |

| | | | | | | | | |
|---|---|---|---|---|---|---|---|---|
| **L686F** | -2.213 | Neutral | 0.13 | TOLERATED | -39 | neutral | 66% | 3/3 |
| **F689V** | -6.474 | Deleterious | 0 | AFFECT PROTEIN FUNCTION | 33 | effect | 66% | 3/3 |
| **L734R** | -3.729 | Deleterious | 0.01 | AFFECT PROTEIN FUNCTION | -17 | neutral | 57% | 2/3 |
| **C746W** | 11.174 | Neutral | 0.02 | AFFECT PROTEIN FUNCTION | -92 | neutral | 97% | 2/3 |
| ***cpdA*** | | | | | | | | |
| **D22A** | -7.763 | Deleterious | 0 | AFFECT PROTEIN FUNCTION | 95 | effect | 95% | 3/3 |
| **H24R** | -7.765 | Deleterious | - | AFFECT PROTEIN FUNCTION | 94 | effect | 95% | 3/3 |
| **H24Y** | -5.825 | Deleterious | 0 | AFFECT PROTEIN FUNCTION | 92 | effect | 95% | 3/3 |
| **S41R** | -4.876 | Deleterious | 0.71 | TOLERATED | 4 | effect | 53% | 2/3 |
| **H165Y** | -5.893 | Deleterious | 0 | AFFECT PROTEIN FUNCTION | 81 | effect | 91% | 3/3 |
| **W174G** | -12.87 | Deleterious | 0.31 | TOLERATED | 82 | effect | 91% | 2/3 |
| **L190Q** | -5.289 | Deleterious | 0.02 | AFFECT PROTEIN FUNCTION | -3 | neutral | 53% | 2/3 |
| **L200R** | -5.507 | Deleterious | 0 | AFFECT PROTEIN FUNCTION | 70 | effect | 85% | 3/3 |
| **G202S** | -5.939 | Deleterious | 0 | AFFECT PROTEIN FUNCTION | 86 | effect | 91% | 3/3 |
| **H203Y** | -5.939 | Deleterious | 0 | AFFECT PROTEIN FUNCTION | 94 | effect | 95% | 3/3 |
| **H205Y** | -5.939 | Deleterious | 0 | AFFECT PROTEIN FUNCTION | 92 | effect | 95% | 3/3 |
| **P221L** | -9.898 | Deleterious | 0 | AFFECT PROTEIN FUNCTION | 87 | effect | 91% | 3/3 |
| **P221T** | -7.919 | Deleterious | 0 | AFFECT PROTEIN FUNCTION | 80 | effect | 91% | 3/3 |
| **S222L** | -5.558 | Deleterious | 0.05 | TOLERATED | 89 | effect | 91% | 2/3 |
| **T255P** | -1.36 | Neutral | 0.2 | TOLERATED | -44 | neutral | 72% | 3/3 |
| ***crp*** | | | | | | | | |
| **L135M** | -1.827 | Neutral | 0 | AFFECT PROTEIN FUNCTION | -52 | neutral | 78% | 2/3 |
| **T141R** | -4.042 | Deleterious | 0.11 | TOLERATED | -37 | neutral | 66% | 2/3 |
| **G142S** | -5.223 | Deleterious | 0.76 | TOLERATED | 50 | effect | 75% | 2/3 |
| **A145T** | -3.522 | Deleterious | - | TOLERATED | -18 | neutral | 57% | 2/3 |
| **A145V** | -3.573 | Deleterious | 1 | TOLERATED | 21 | effect | 63% | 2/3 |
| **L149R** | -4.97 | Deleterious | 0.03 | AFFECT PROTEIN FUNCTION | 42 | effect | 71% | 3/3 |
| **M190R** | -0.079 | Neutral | 0.47 | TOLERATED | -49 | neutral | 72% | 3/3 |
| **L196R** | -5.087 | Deleterious | 0 | AFFECT PROTEIN FUNCTION | 67 | effect | 80% | 3/3 |

[a] frame-shift mutations were not investigated here
[b] PROVEAN [22]: variants with a score equal to or below -2.5 are considered "deleterious"; variants with a score above -2.5 are considered "neutral"
[c] SIFT [23]: amino acids with probabilities < 0.05 are predicted to be deleterious
[d] SNAP2 [24]: values range from -100 (neutral) to 100 (effect)
[e] red: deleterious effect on protein function; green: neutral effect on protein function

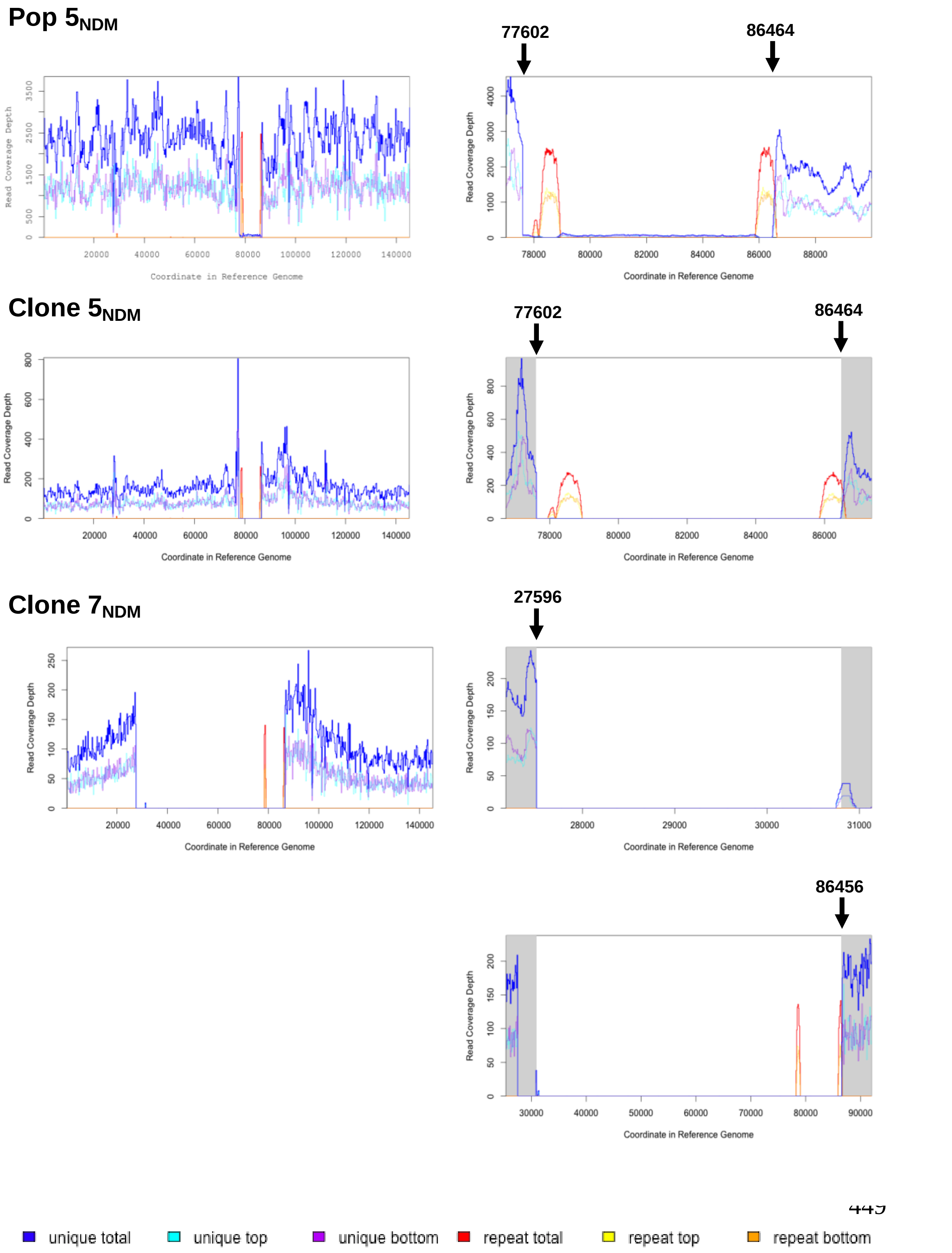


**Supplementary Figure 1.** Read coverage plots of evolved pK71-77-1-NDM in evolved Pop $5_{NDM}$, Clone $5_{NDM}$ and Clone $7_{NDM}$. Large deletions in the evolved plasmid sequences were identified when mapping short-read sequencing data against the ancestral pK71-77-1-NDM using breseq [10] (see also main text Methods). Plots were part of the breseq output. The read coverage depth across the whole plasmid (145 kb; left column) and around specified areas with missing coverage (white background; right column) is shown. Beginning and end positions of the largest possible deletion (including repetitive regions indicated in red/yellow/orange) are displayed and were accessed using the bam2cov command in breseq. A colour legend common to all plots is given at the bottom.

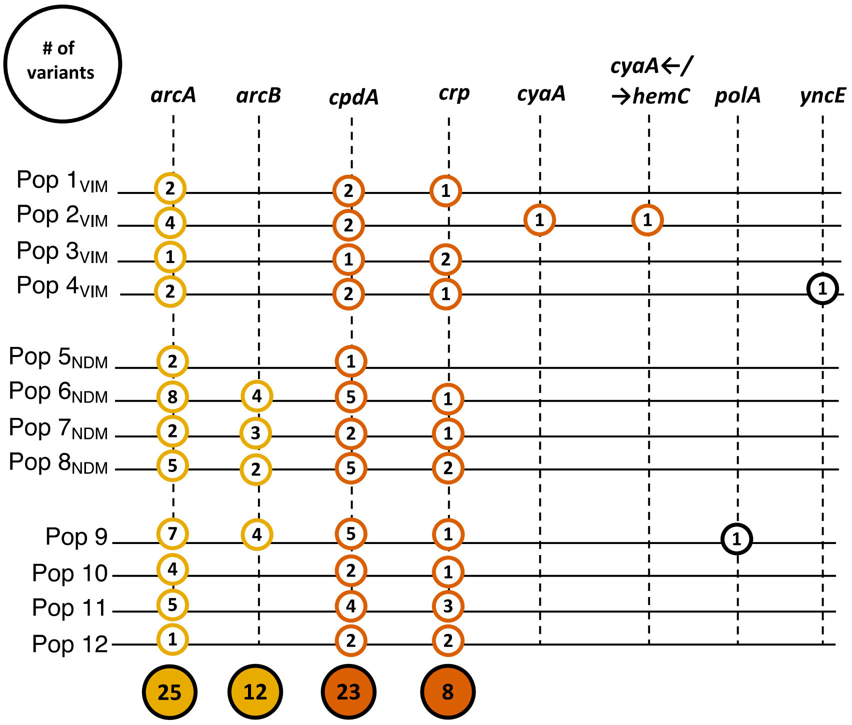


**Supplementary Figure 2.** Number of chromosomal variants per evolved population and mutational target gene. Total number of unique variants per gene are summarized in filled circles.

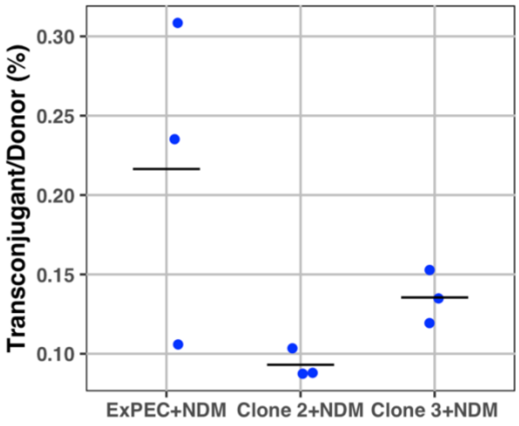


**Supplementary Figure 3.** Conjugation frequency of ancestral pK71-77-1-NDM from ancestral and evolved backgrounds (ExPEC+NDM and Clone 2+NDM/Clone 3+NDM, respectively) into strain K56-43 RifR. The plasmid was transferred with an average frequency of 0.22 ± 0.10% from strain ExPEC+NDM, 0.09 ± 0.01% from Clone 2+NDM and 0.13 ± 0.02% from Clone 3+NDM (all n = 3).

**Supplementary Figures 4 and 5.** Ontology graphs of enrichment and overrepresentation analysis RNA-Seq. Up and downregulated processes (Supp.Figure 4 and 5, respectively) in Clone 2 and 3 with and without pG06-VIM-1 due to adaptive chromosomal mutations. Border colour indicates significant changes for: Clone 2+VIM vs. ExPEC+VIM (dark blue); Clone 3+VIM vs. ExPEC+VIM (red); both Clone 2+VIM and 3+VIM vs. ExPEC+VIM (yellow). Fill colour indicates significant changes for: Clone 2 vs. ExPEC (light blue); Clone 3 vs. ExPEC (purple); both Clone 2 and 3 vs. ExPEC (grey). The graph is read from the bottom, such that a lower biological process “is a“ (black lines) or is “part of” (light blue lines) a higher connecting process.

**Supplementary Figure 6.** Ontology graph of enrichment and overrepresentation analysis RNA-Seq. Downregulated processes due to pG06-VIM-1 acquisition in Clone 2+VIM vs. ExPEC+VIM. Fill colour: significant changes in Clone 2+VIM (light blue). The graph is read from the bottom, such that a lower biological process “is a“ (black lines) or is “part of” (light blue lines) a higher connecting process.
