## Supplementary figures and images for "Piggybacking on niche-adaptation reduces the cost of multidrug resistance plasmids"

### Supplementary Figure 4

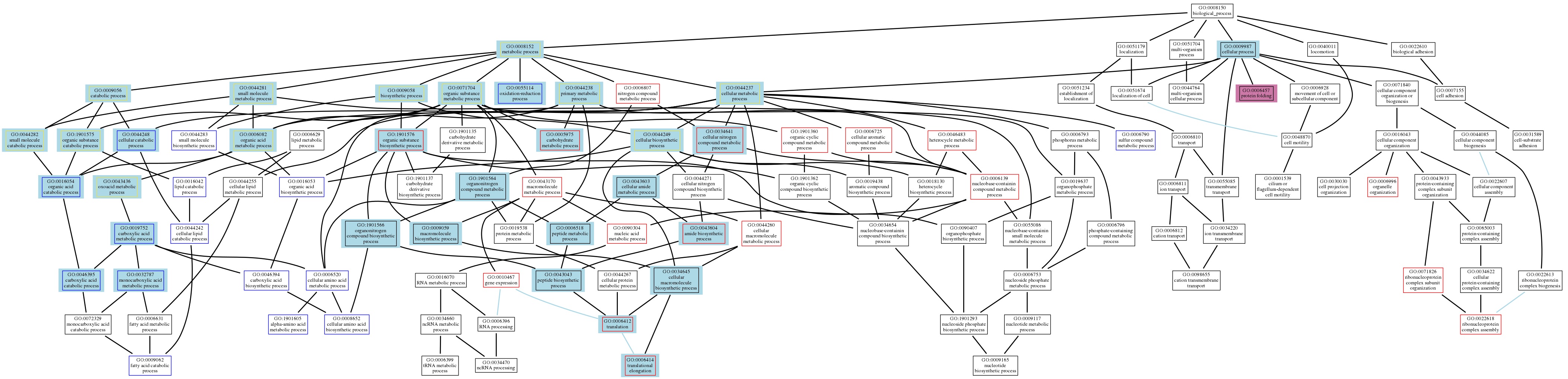

### Supplementary Figure 5

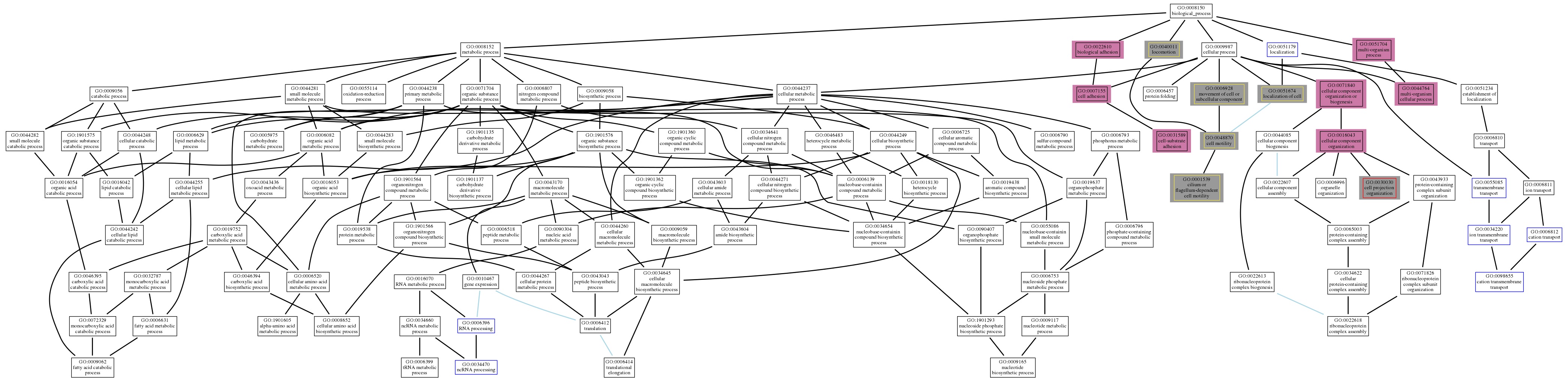

### Supplementary Figure 6

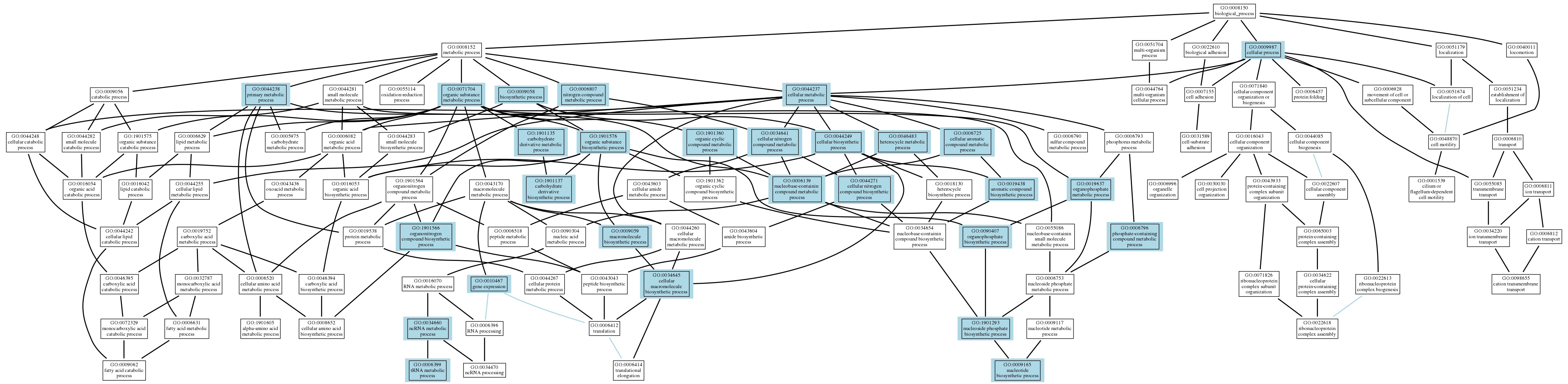
